# A small molecule that promotes cellular senescence prevents fibrogenesis and tumorigenesis *in vitro*

**DOI:** 10.1101/2021.06.01.446522

**Authors:** Moon Kee Meang, Saesbyeol Kim, Ik-Hwan Kim, Han-Soo Kim, Byung-Soo Youn

## Abstract

**Background:** Uncontrolled proliferative diseases such as fibrosis or cancer can be fatal. We previously found that a compound containing a chromone scaffold, ONG41008, had potent anti-fibrotic effects in diseased human lung myofibroblasts but not normal human lung fibroblasts.

**Methods:** We investigated the effects of ONG41008 on tumor cells, and compared these effects with those in pathogenic myofibrotic cells and normal fibroblasts cells.

**Findings:** Stimulation of A549 lung carcinoma epithelial cells with ONG41008 resulted in cellular senescence, indicating that dysregulated cell proliferation is common to fibrotic cells and tumor cells. Replicative senescence of A549 cells resulted in multinucleation, which was followed by oncogene-induced senescence. There was significant upregulation of expression and nuclear translocation of p-TP53 and p16 in ONG41008-treated A549 cells, and all cells died after 72 hr. Similar effects occurred after ONG41008 treatment in several human aggressive cancer cell lines such as PANC1, MCF7, PC3, or primary non-small cell lung carcinoma cells. Unlike cisplatin, ONG41008 was not toxic to normal human lung fibroblasts or primary prostate epithelial cells, suggesting ONG41008 can distinguish the intracellular microenvironment between normal cells and aged or diseased cells. This effect might occur as a result of the increased NAD/NADH ratio or increased lactate dehydrogenase levels in aged or diseased cells.

**Interpretation:** To our best knowledge, this is the first study to show that a small molecule can arrest uncontrolled proliferation during fibrogenesis or tumorigenesis *in vitro*. ONG41008 could be a potential drug for a broad range of fibrotic or tumorigenic diseases.

**Research in Context:** *Evidence before this study:* The notion that aging is a disease and that diseases occur as a consequence of aging was first put forward by David Sinclair and colleagues (*Aging Cell*;14(4):497-510). There are biological systems that provide evidence for this notion; for example, somatic cells can revert to embryonic cells, producing younger somatic cells. This phenomenon underlies induced pluripotent stem cells. Another example is that some types of jellyfish can live forever. These examples suggest that a counter-aging program exists in animals. Human diseases are the manifestations of cell aging generated by the accumulation of somatic mutations. Aged and pathogenic cells are senescent, so a drug that specifically targeted senescent cells might initiate a cellular program that could ameliorate age-associated disease. Indeed, the kinase inhibitor dasatinib induces cellular senescence (*Clin Ther* 2007 29:2289-2308). In 2017, two drugs that target senescent cells were identified: quercetin and fisetin. These drugs selectively kill senescent cells, and are referred to as senolytic drugs or senotherapeutics (*Aging* 2017 8;9(3):955-963). Although it is established that senescent cells accumulate in cancer and idiopathic pulmonary fibrosis (*Nat Commun* 2017 23;8 :14532), the effect of senolytic drugs in these diseases is largely unknown.

*Added value of this study:* This study characterized a novel drug, termed ONG41008, which was found to have both senogenic and senolytic effects in cell-based assays. ONG41008 induced senescence in myofibroblasts and several cancer cell lines representative of aggressive human cancers, which was followed by cell death. Importantly, ONG41008 exhibited essentially no toxicity on normal human lung fibroblasts or primary prostate epithelial cells.

*Implication of all the evidence:* Based on our results, we believe that ONG41008 is a potent inducer of cellular senescence (replicative senescence and oncogene-induced senescence) and causes arrest of uncontrolled, pathogenic proliferation of myofibroblasts or cancer cells.

## Introduction

Uncontrolled progression of the cell cycle is linked to the development of fibrosis and cancer (1). The entry of cells into the cell cycle and exit from the cell cycle needs to be tightly regulated (2), otherwise tumorigenesis can occur (3). Although targeting cell-cycle progression using small molecules is a potential anti-cancer strategy, it may be associated with undesirable risks such as elimination of bystander non-cancerous cells, systemic inhibition of stem-cell differentiation or aggravation of homeostatic immunity (4). Compounds that contain chemical scaffolds found in natural products, such as the chromone scaffold (CS) found largely in flavones or isoflavones, might be useful anti-proliferative drugs; these natural products are largely safe and suppress uncontrolled proliferation, as shown in studies using pathogenic myofibroblasts or aggressive tumor cells (5). Metastasis plays a key role in the additional development of cancer, and so targeting metastasis is an active area of research (6). It is increasingly clear that metabolic regulation in cancer cells is closely coupled to the progression of the cell cycle. Moreover, it is well-established that tumor-cell growth is exclusively dependent on aerobic glycolysis (also known as lactate fermentation) (7). Similarly, pathogenic myofibroblasts utilize aerobic glycolysis and tend to generate a hypoxic environment, so aerobic glycolysis also plays a central role in the initiation and perpetuation of fibrosis (8).

Flavones are members of the polyphenol family of compounds, which contains over 10,000 compounds exclusively found in the plant kingdom (9). In general, these phytochemicals protect plants from radiation damage (10). Due to their anti-oxidant or anti-inflammatory potential, flavones have long been used to treat inflammatory diseases such as arthritis and asthma (11). Chromone, 1,4-benzopyrone-4-one, is a central chemical scaffold that is found in flavones and isoflavones (12). We recently reported that eupatilin, a CS-containing compound isolated from an *Artemisia* species and its synthetic analog ONG41008 inhibit fibrogenesis *in vitro* and fibrosis *in vivo.* These effects occurred as a result of actin depolymerization followed by disassembly of the latent TGFβ complex (LTC), resulting in inhibition of epithelial–mesenchymal transition (EMT) (13). Here, we show that ONG41008 potently induces replicative senescence (RS) in pathogenic myofibroblasts and oncogene-induced senescence (OIS) in several cancer cell lines being representative of aggressive forms of human cancer, resulting in cellular escape from fibrogenesis and tumorigenesis. These anti-proliferative actions of ONG41008 suggest it could be a new therapeutic modality for treating fibrosis as well as cancer.

## Results

### ONG41008 causes cell-growth arrest and replicative senescence (RS) in DHLF

Diseased human lung fibroblasts (DHLF) from idiopathy pulmonary fibrosis (IPF) patients are αSMA+ pathogenic myofibroblasts that express a range of muscular collagens. We sought to identify if the mechanism of action of ONG41008 was different from that of two first-in-class IPF drugs: nintedanib and pirfenidone. Nintedanib, which has high cellular toxicity, was originally developed as an anti-cancer drug and has been repurposed as an anti-IPF drug. Pirfenidone is hepatotoxic at high concentrations and indirectly modulates the function of immune cells, resulting in reduced activity of TGFβ (14) (15). We recently reported that ONG41008 inhibited TGFβ biogenesis, thereby blocking TGFR signaling and reprogramming the EMT (13). In the current study, as shown in Figure 1A, nintedanib induced robust cell death in DHLF at 24 hr, and after a longer exposure of 48 hr, nearly all DHLF were dead. The IC_50_ value of nintedanib was 10.17 μM at 72 hr. Pirfenidone had no effect on the survival of DHLF. Interestingly, ONG41008 partially affected cell survival; the cell survival rate was more than 60% even when concentrations of ONG41008 were increased, meaning we were unable to obtain an IC_50_ value. This result strongly indicates that ONG41008 treatment caused systemic growth arrest of DHLF. As a control experiment, normal human lung fibroblasts (NHLF) were treated with nintedanib, pirfenidone, and ONG41008. Surprisingly, although nintedanib highly inhibited cell survival (the IC_50_ value was around 30 μM; Figure 1B) and pirfenidone did not affect cell survival, ONG41008 did not affect the survival of NHLF. This finding meant we were not able to calculate an IC_50_ value for ONG41008. This series of experiments showed that DHLF were more susceptible than NHLF to the pro-apoptotic effects of nintedanib.

**Figure 1.**
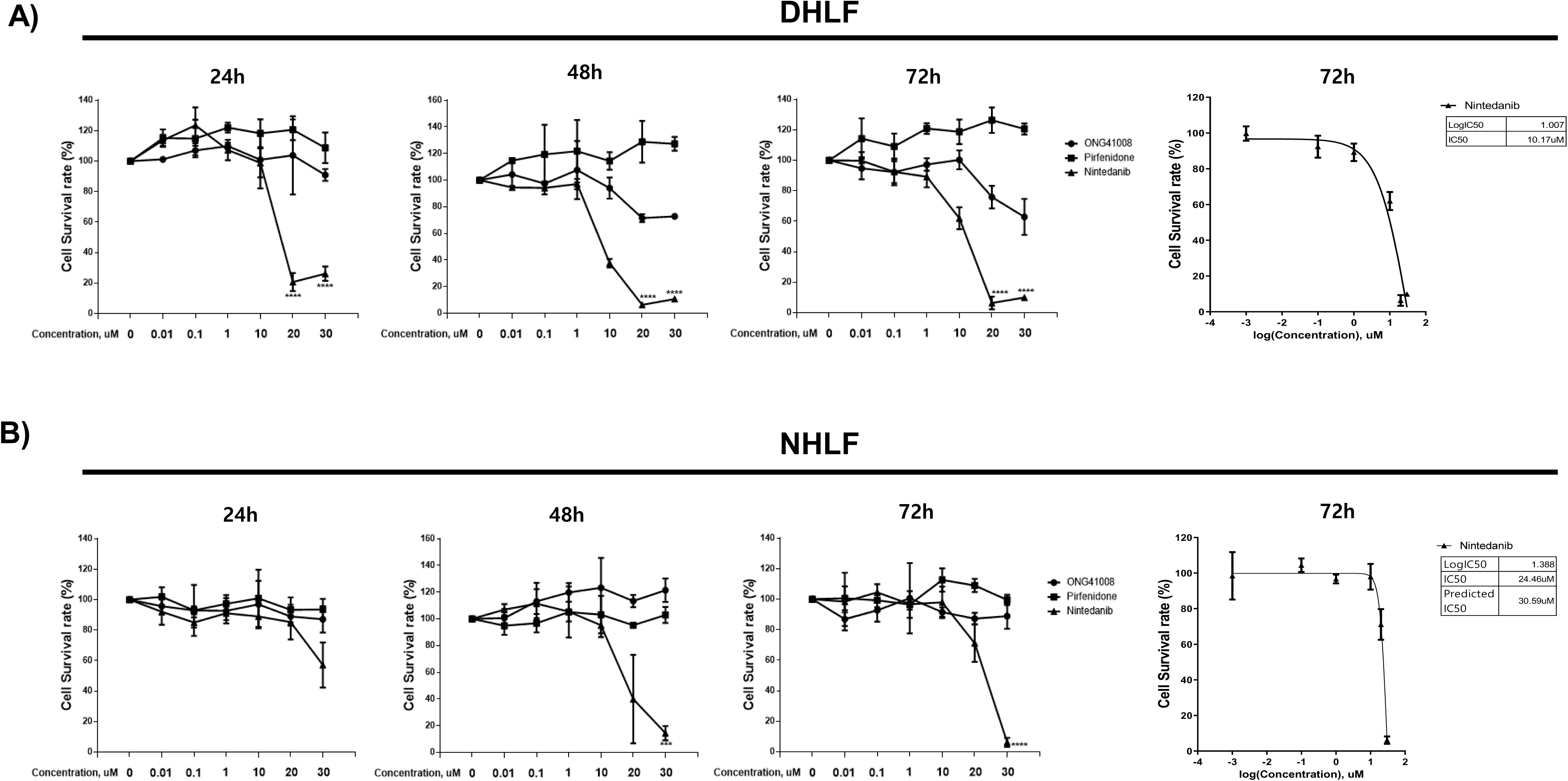
Comparison of the survival rate of DHLF and NHLF treated with ONG41008, nintedanib, or pirfenidone. A-B) DHLF and NHLF were stimulated with various concentrations (up to 30 μM) of the indicated drugs for 24 hr, 48 hr, and 72 hr. The cell survival rate was measured using a CCK-8 assay. The IC_50_ value of nintedanib was calculated with a sigmoidal, four-parameter logistic in GraphPad Prism version 7.00. At least three independent measurements were conducted.

To rule out the involvement of apoptosis in ONG41008-mediated growth arrest in DHLF, three apoptotic features were measured using: 1) an activated caspase-3 assay, 2) measurement of the mitochondrial membrane potential (MTMP), and 3) an LDH cytotoxicity assay. FCCP (trifluoromethoxy carbonylcyanide phenylhydrazone) was used as a control in MTMP experiements. As shown in Figures 2A, 2B, and 2C, only nintedanib caused significant apoptosis of DHLF in the three assays, allowing estimation of respective IC_50_ or EC_50_ values. ONG41008 and pirfenidone were unable to elicit pro-apoptotic pathways. This result suggests that the systemic growth arrest induced by ONG41008 was not a consequence of apoptosis and instead is likely related to cellular senescence.

**Figure 2.**
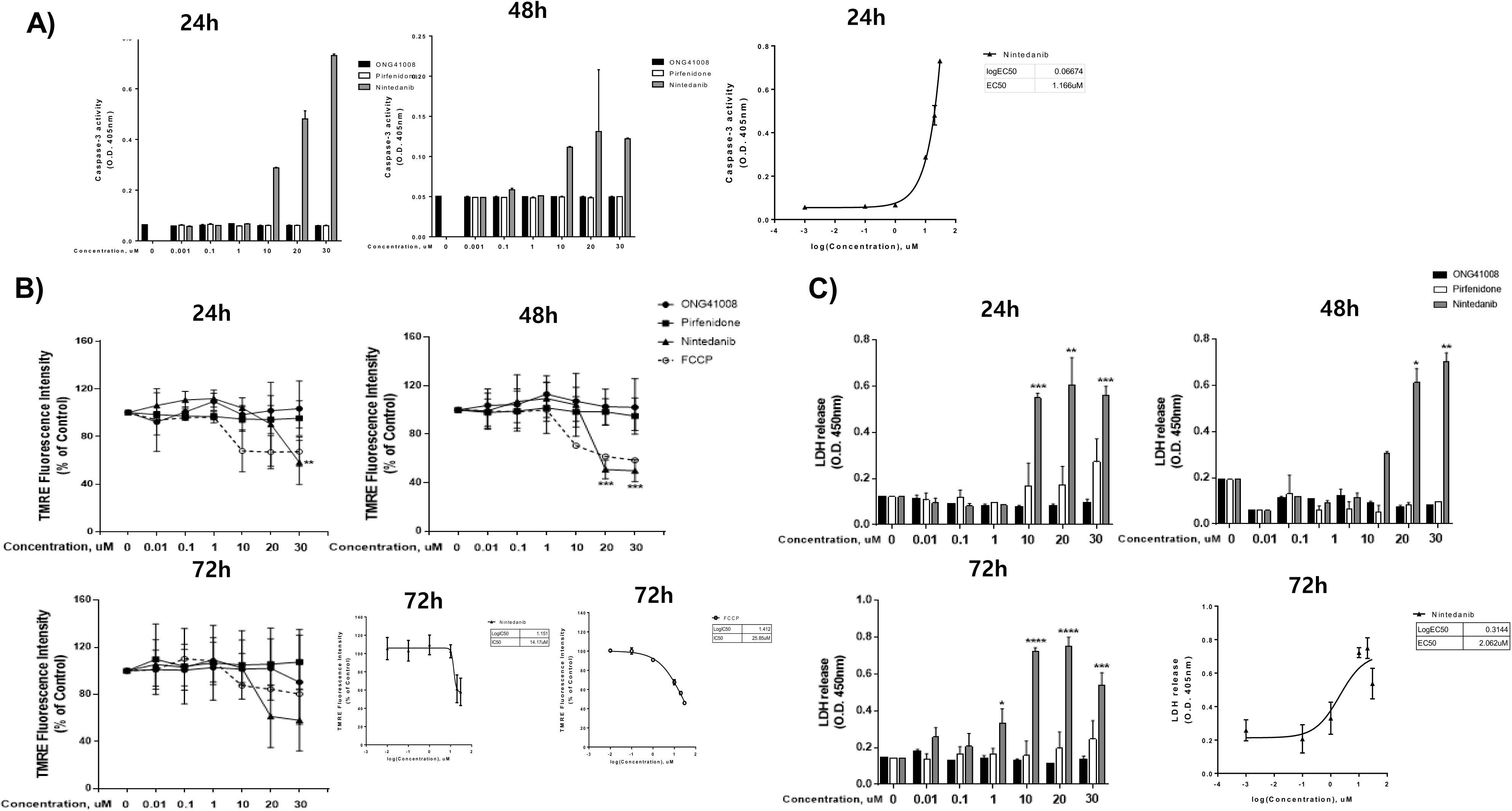
ONG41008 did not induce apoptosis in DHLF. DHLF were stimulated with ONG41008, pirfenidone, or nintedanib. A) An activated caspase-3 assay was performed and the EC_50_ value of nintedanib was calculated at 24 hr. B) The mitochondrial membrane potential was measured. The mitochondrial oxidative phosphorylation uncoupler FCCP was used as a control. The IC_50_ values of nintedanib and FCCP were calculated at 72 hr. C) Lactate dehydrogenase (LDH) levels were assayed. The EC_50_ value was calculated with a sigmoidal, four-parameter logistic in GraphPad Prism version 7.00. At least three independent measurements were conducted.

### ONG41008 is a potent senogenic molecule

Senogenic and senolytic compounds were first identified in 2017 (16) (17) (18). Dasatinib, a Bcr-Abl and Src kinase inhibitor, was the first clinically used drug that induces cellular senescence. Quercetin and fisetin were pioneering senolytic drugs (19) (20). Interestingly, like quercetin and fisetin, ONG41008 contains a CS (21). We previously reported that three compounds containing a CS had potent anti-fibrogenic activity (13), of which ONG41008 had the highest anti-fibrogenic capability. An immunocytochemistry (ICC) experiment showed that robust cellular senescence occurred when A549 cells were stimulated with ONG41008 and MNC were clearly formed (Supplementary Figure 1). MNC is an early signature for mitotic slippage followed by senolysis (22). It has been well documented that the cellular senescence pertinent to cancer cells refers to oncogene-induced senescence (OIS) (23). Figure 3 summarizes anti-fibrogenic, senogenic, and senolytic properties associated with the chromone scaffold-containing chemical structures. It has been shown previously that the presence of a methoxy group at the C6 position in the CS is essential to mediate anti-fibrogenic effects, and as such, apigenin, quercetin, and fisetin were not anti-fibrogenic. ONG41008 was anti-fibrogenic, senogenic, and senolytic. However, chemical analogs of ONG41008, namely hispidulin, jaceosidin, and apigenin did not induce cellular senescence.

**Figure 3.**
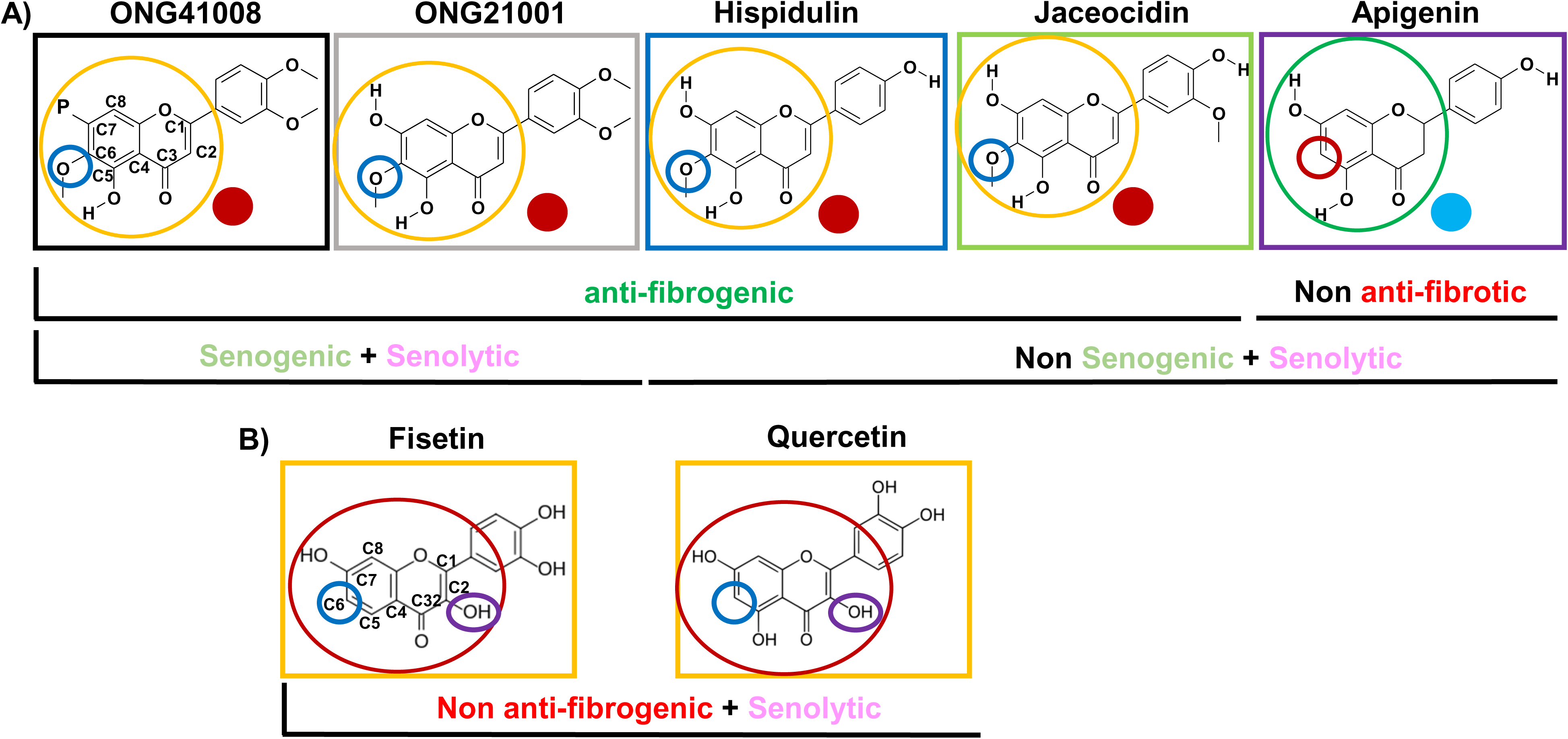
The chemical structure of compounds containing a chromone scaffold. ONG41008, ONG21001, hispidulin, and jaceosidin, which are denoted by red circles, have anti-fibrogenicc effects and Apigenin is not anti-fibrogenic, demonstrated by a blue circle. The presence of a methoxy group at C6 in the CS is necessary for anti-fibrogenic capacity ONG41008 and ONG21001 have senogenic and senolytic effects. No anti-fibrogenic capacities and senogenic activity are associated with the structure’s quercetin and fisetin.

A hallmark of RS or OIS is cell flatness (24). We previously found that GATA6 plays an important role in perpetuating EMT in DHLF (13). We assumed that the expression of GATA6 would be reduced in the absence of TGFβ. We used time-lapse microscopy to monitor the effects of increasing concentrations of ONG41008 in an RS model in DHLF. As shown in Figure 4A, GATA6 expression was consistently observed throughout the study; the reasons for this finding remain to be elucidated. RS occurred at 5 μM ONG41008, became more evident at 10 μM ONG41008 and the maximal effect on RS occurred at 20 μM ONG41008. H2AX is a senescence-specific histone. (25). To further substantiate ONG41008-mediated RS, DHLF were stimulated with 20μM ONG41008 for 24h, 48h, or 72h and stained with anti-H2AX. As shown in Figure 4B, H2AX were nuclear localized and heavily stained, suggesting that ONG41008 is a potent inducer of RS. In the absence of ONG41008, DHLF did not undergo RS. Video analysis also showed that 10 μM ONG41008 strongly induced RS (Supplementary Video I). To compare senogenic capability of ONG41008 with that of dasatinib in conjunction with quercetin or fisetin DHLF were stimulated with these drugs including pirfenidone, which is one of two first-in class IPF drugs as control and ICC was conducted with the use of anti-p16. As seen in Figure 4C, ONG41008 induced a robust RS and fisetin and dasatinib weakly induced RS in DHLF. Quercetin and pirfenidone were not able to induce RS.

**Figure 4.**
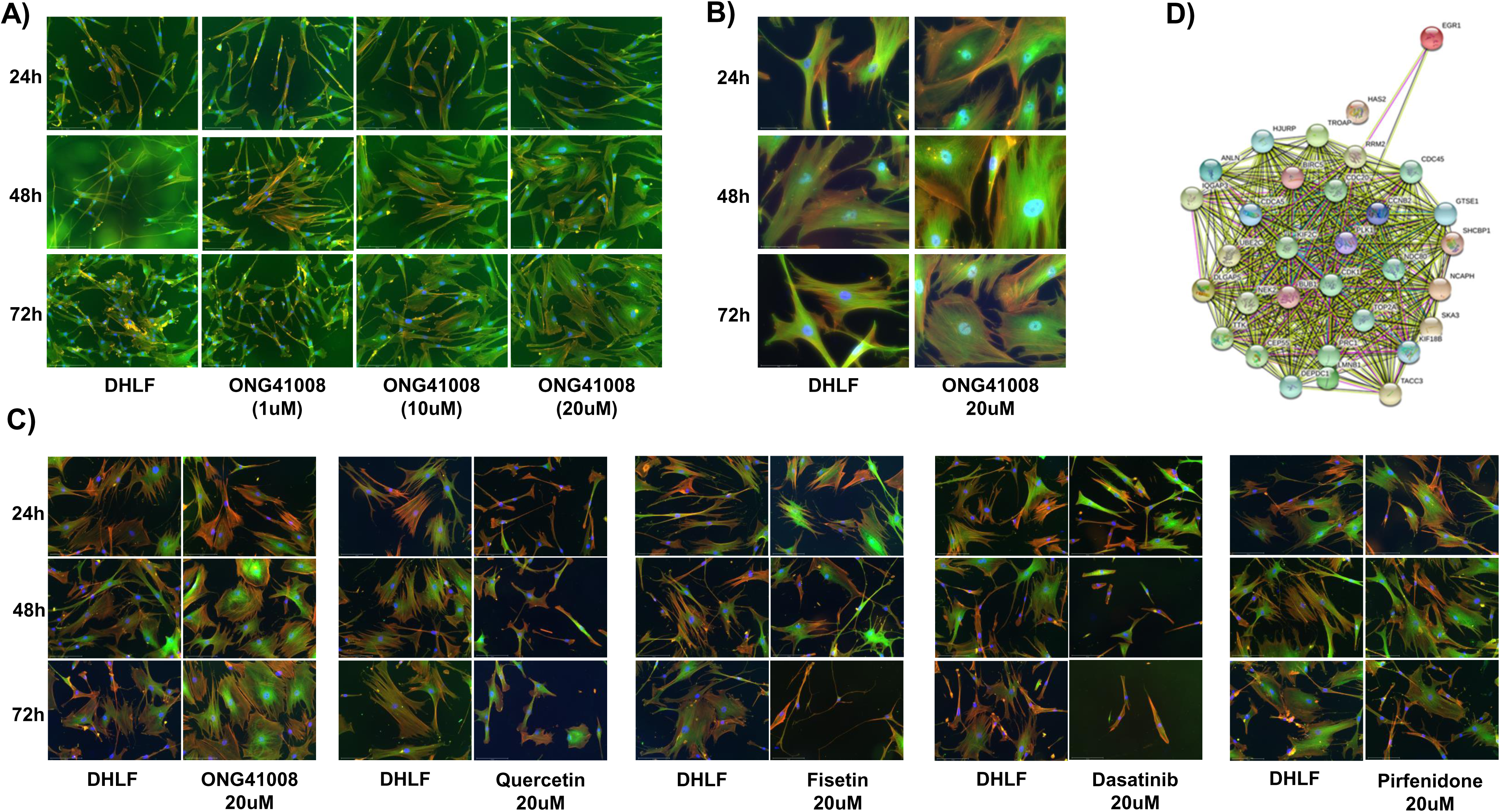
Induction of RS by ONG41008 and an interactome analysis. A) DHLF were treated with different concentrations of ONG41008 or a control (RPMI medium). Immunocytochemistry (ICC) was conducted with anti-human GATA6, phalloidin, and DAPI. Morphological changes were monitored under phase-contrast and fluorescence microscopy. Images that show typical replicative senescent cells are denoted by the phrase “replicative senescence”. B) DHLF were stimulated for 24h, 48h, or 72h and stained with anti-H2AX. ICC images were mounted. C) An RNA-seq analysis was conducted using total RNAs prepared from untreated DHLF or 10 μM ONG41008-treated DHLF. An interactome representing upregulated genes was established using the STRING program, based upon p-values (>0.005). D) DHLF were stimulated with 20μM ONG41008, 20μM Quercetin, 20μM Fisetin, 20μM Dasatinib, or 20μM Pirfenidone for 24h, 48h, or 72h and replicative senescence was compared via ICC with the use of anti-p16.

Next, we conducted an RNA-seq analysis, giving rise to the nuclear interactome shown in Figure 3B. EGR1 (early growth response 1) seemed to be a priming protein, which interacted with proteins including CDC45 (cell division cycle 45), TACC3 (transforming acidic coiled-coil-containing protein 3 and CDK1 (cyclin-dependent kinase 1). These proteins play pivotal roles in the onset of nuclear reprogramming for trans-cell differentiation, cell-cycle control, and cellular senescence (26) (27) (28) (29). We also discovered that unique transcriptomes are responsible for driving RS; RNA-seq analysis was conducted to compare differences in gene expression between untreated DHLF and 10 μM ONG41008-treated DHLF. Upregulated genes were sorted according to their p-values. As shown in Supplementary Figure 2 and Supplementary Figure 3, three interactomes seemed to be working cooperatively when cells undergo RS; the first was a metabolic interactome typified by pyruvate dehydrogenase kinase (PDK)1, the second was actin biogenesis, and the last was related to histone modification, DNA replication, and the generation of a muscle–neuron signature. The reason why a muscle–neuron signature is generated for RS remains to be explored. Interestingly, the vast majority of top-ranked genes affected by ONG41008 were soluble factors or receptors (Supplementary Table I). We speculate that to drive RS, expression of these cytokines or their interaction with cognate receptors is be required to prime cellular senescence prior to the formation of euchromatin formation.

Taken together, ONG41008 is a potent inducer of cellular senescence in pathogenic myofibroblasts and cancer cells and contains CS but are distinguished from quercetin and fisetin, which are exclusively senolytic molecules in cancer cells.

### ONG41008 is senolytic in cancer cells

A549 cells were stimulated with ONG41008, quercetin, fisetin, or dasatinib (as a senogenic control). ONG41008, quercetin, and fisetin induced senolysis that was characterized by the MNC formation (Supplementary Figure 4, highlighted by white circles). Dasatinib was toxic to A549 cells. In particular, A549 became rapidly senescent when treated with 10 μM ONG41008. Although fisetin appeared to induce senescence, but when treated with 20 μM quercetin or fisetin, the flat cell morphology disappeared and even appeared shrunken under higher magnification. No MNC was observed in DHLF stimulated with ONG41008 and we were unable to confirm senolysis of DHLF stimulated with ONG41008, which remains to further be elucidated. To quantitate the senolytic activity of ONG41008, quercetin, and fisetin containing a CS, CCK-8 assays were conducted by using PANC1 cells and the IC_50_ values of these compounds were compared as shown in Figure 5A. Cisplatin was used as an apoptosis control. Quercetin exhibited the highest senolytic activity, and ONG41008 and fisetin also showed significant senolytic activity. The finding that quercetin and fisetin showed senolytic activity indicates that senescence may have occurred in PANC1 cells. Caspase3,7 assay is a good tool for determining late stage senolysis, apoptosis. Interestingly, ONG41008 only exhibited a decent senolytic activity in terms of activation of caspase3,7 in conjunction with dasatinib and cisplatin (Figure 5B). Further studies are needed to determine the effects of these compounds on cellular senescence and senolysis in more cancer cell lines and human primary cells. In addition to A549 cells, ONG41008-mediated OIS was explored in malignant human cancer cell lines; the human triple-negative (TNBC) breast cancer cell line MCF7, the human pancreatic ductal carcinoma cell line PANC1, and PC3 cells (an MDR+ / multidrug-resistant human prostate cancer cell line). These cell lines were stimulated with ONG41008 for 48 hr and ICC was conducted. All four cancer cell lines showed robust cellular senescence, as shown in Supplementary Figure 5. MNC is denoted by white circles and presumable mitosis in a proportion of A549 cells is represented by green circles.

**Figure 5.**
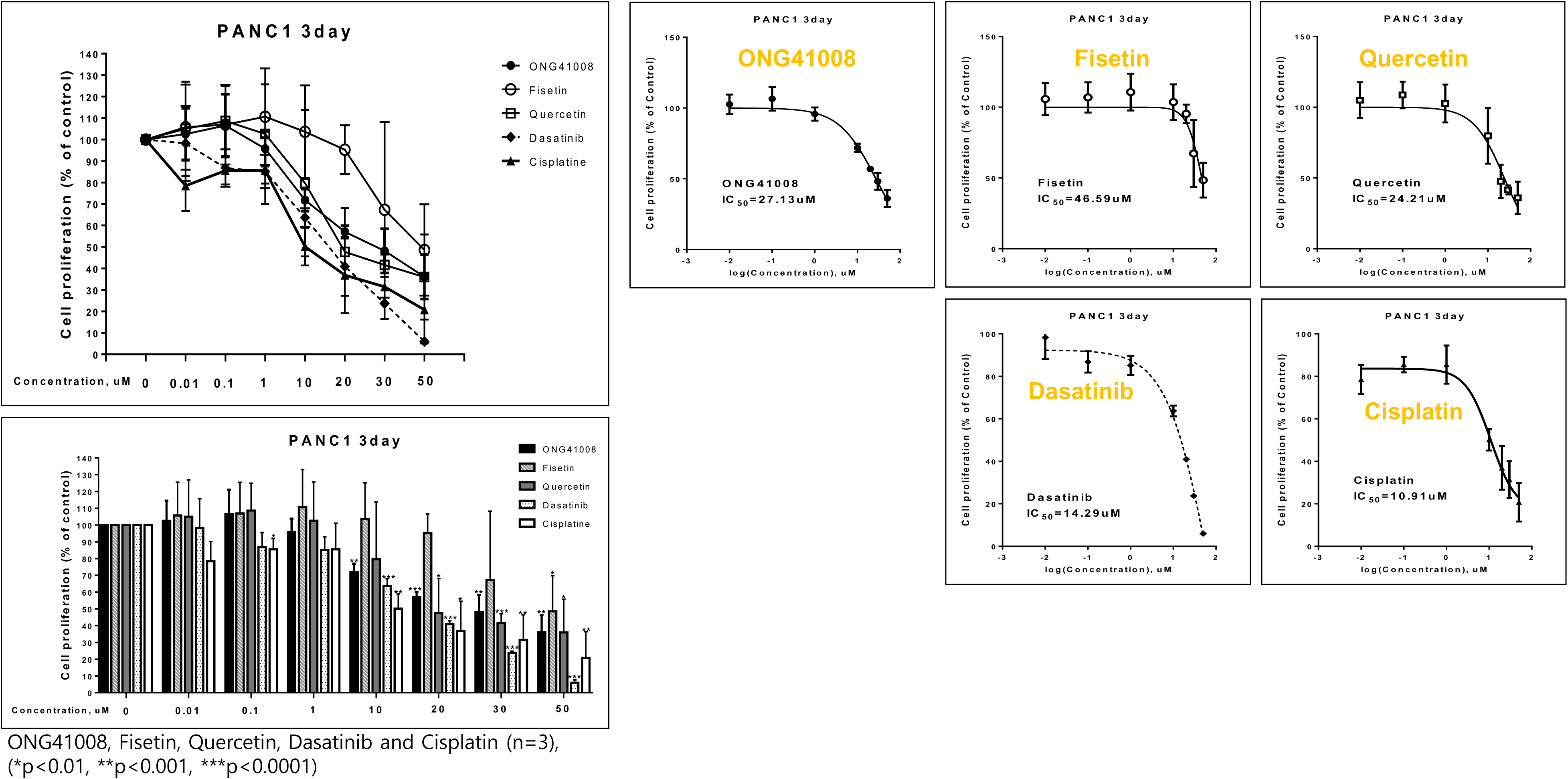
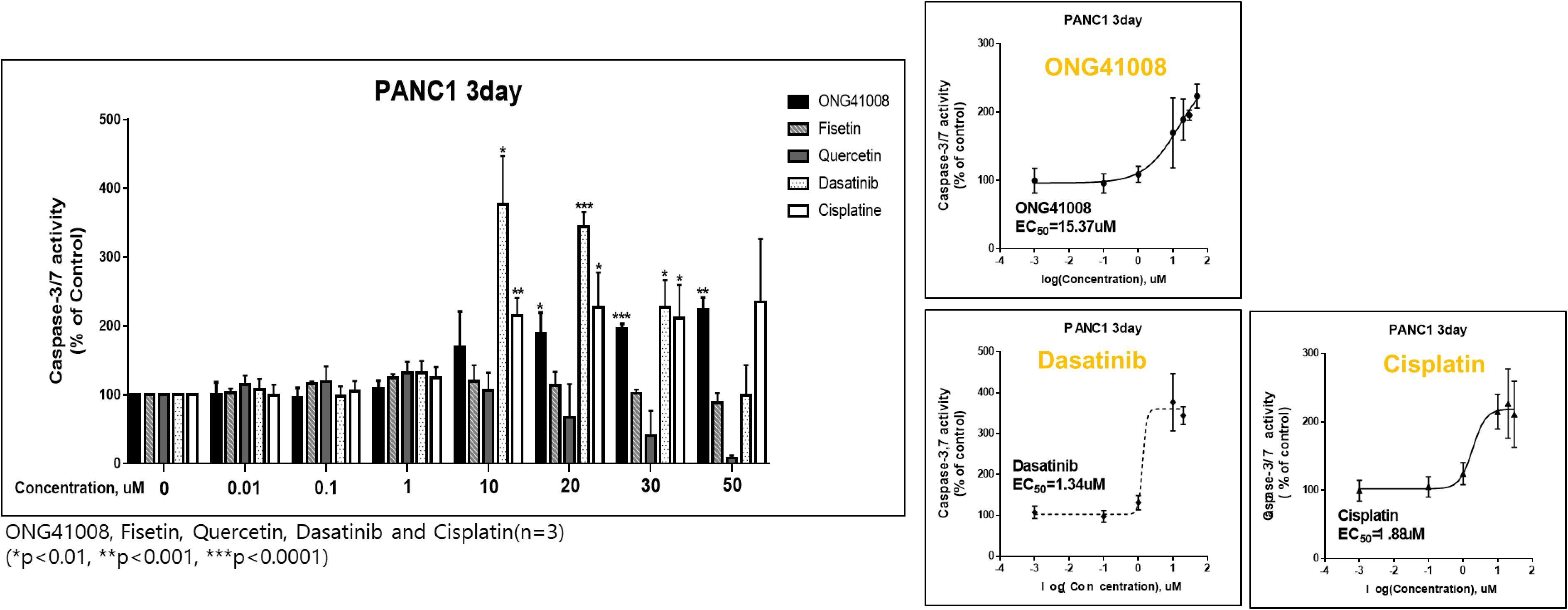
Cellular senescence and the senolytic capability of quercetin, fisetin, ONG41008 and dasatinib. A) A549 were treated with increasing concentrations (100nM from to 50μM) of ONG41008, Quercetin, Fisetin, Dasatinib, or Cisplatin. CCK-8 assays were conducted. IC_50_ values were calculated. B) A549 were stimulated as described above and Caspase3,7 assays were conducted. EC_50_ values were also calculated. Three independent experiments with triplicates were performed and statistical acquisition was completed ONG41008, Fisetin, Quercetin, and Cisplatin (*p<0.01, **p<0.001, ***p<0.0001)

Taken together, these results show that ONG41008 is both senogenic and senolytic as well as previously anti-fibrogenic, suggesting that the therapeutic potential of ONG41008 could be extended to fibrosis as well as cancer.

### ONG41008-mediated RS of DHLF requires TP53, p21 and p16 activity

Several tumor suppressor proteins are responsible for halting the progression of the cell cycle (30). TP53 is phosphorylated and couples mainly to p21 or p16, leading to cell-cycle inhibition. As shown in Figure 6A, 1 μM ONG41008 treatment did not alter the expression of TP53 but caused rapid translocation of this to the nucleus 1 hr after treatment. Treatment of DHLF with 1 to 10 μM ONG41008 upregulated p21 expression and caused translocation of the protein to the nucleus (Figure 6B). The expression of p16 was also upregulated and translocated to both the perinuclear zone and the nucleus (Figure 6C).

**Figure 6.**
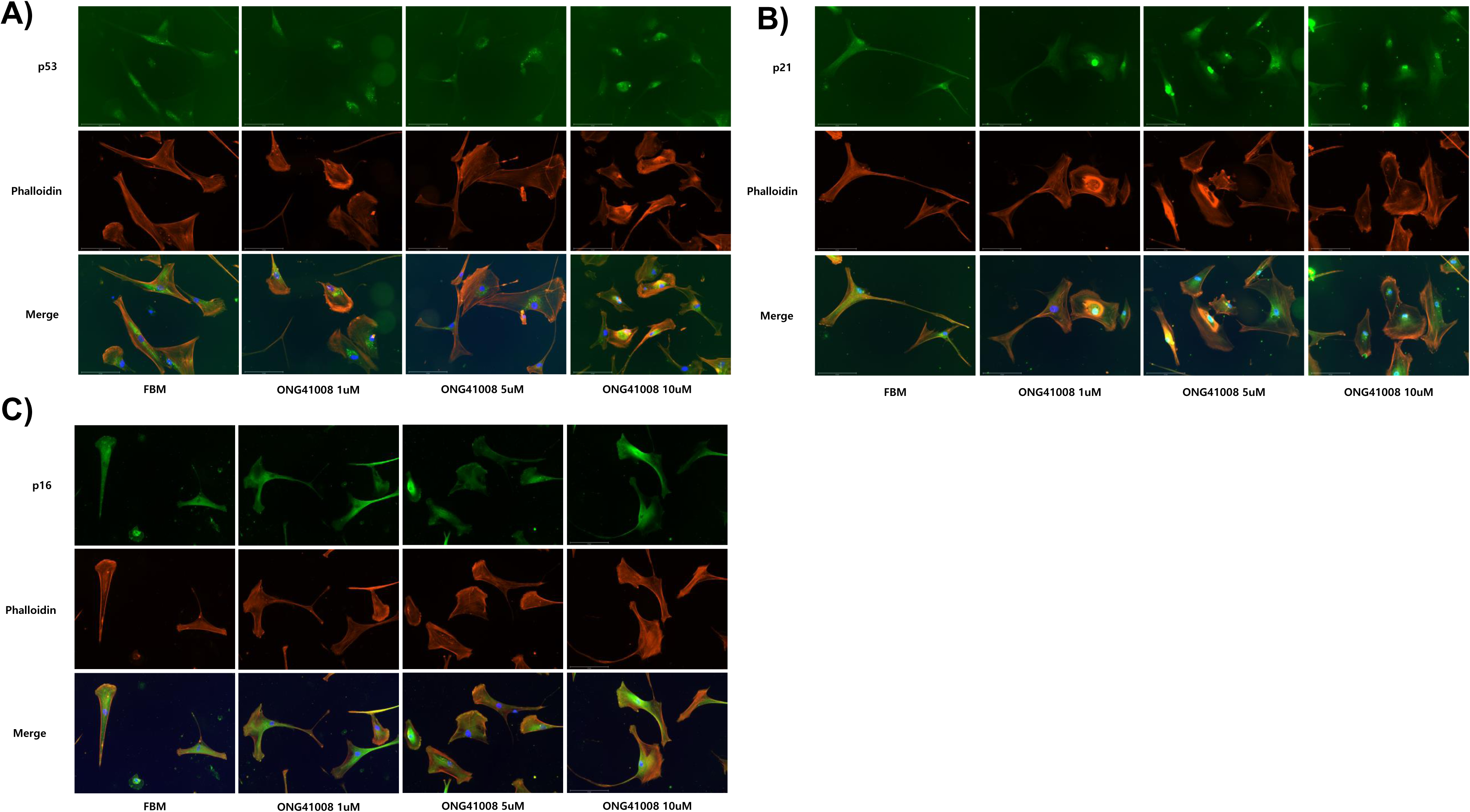
Translocation of TP53 to the nucleus, and induction and nuclear translocation of p21 and p16 in DHLF upon ONG41008 treatment. DHLF were treated with 1 μM to 10 μM ONG41008 or a control (RPMI medium). ICC was conducted with anti-human TP53, anti-p21, anti-p16, or phalloidin in conjunction with DAPI. A) Translocation of p53 to the nucleus. B) Induction and translocation of p21 to the nucleus. C) Induction and translocation of p16 to the nucleus.

### Biochemical analysis of ONG41008-mediated OIS in A549 and PANC1

Pathogenic myofibroblasts and tumor cells share several features; first, aerobic glycolysis is the major pathway for the catabolism of glucose. Second, uncontrolled cell division occurs. Third, their survival is dependent on immune escape. The observation that ONG41008 rapidly caused DHLF to become RS prompted us to see if cancer cells would behave similarly to DHLF. Cellular senescence was established in A549 cells by treatment with 10 μM ONG41008 for 24hr, 48hr, or 72 hr. Similar to our studies of RS in DHLF, we explored changes in the expression and location of TP53, p21, and p16, which can induce OIS in A549 cells. As shown in Supplementary Figure 6A, TP53 protein expression remained unchanged during ONG41008 treatment, but 1 μM ONG4008 caused nuclear translocation of TP53. Expression of p21 was upregulated and similarly localized in the nucleus (Supplementary Figure 6B). Expression of p16 was concentration-dependently upregulated by ONG41008, and was localized in the nucleus and distributed at the perinuclear zones (Supplementary Figure 6C). We discovered that cell morphology was more homogenous 72hr after ONG41008 treatment than 24hr after treatment; this result may reflect the elimination of multinucleated A549 cells. Moreover, maximal translocation of p16 to the nucleus was complete at 72 hr. We anticipate that p16-saturated A549 cells would undergo apoptosis (Supplementary Figure 7). Since the ability of TP53 to regulate cell-cycle arrest is dependent on its phosphorylation of (31), we conducted western blot analysis using a phospho-specific TP53 antibody. As shown in Figure 7A, ONG41008 concentration-dependently increased the level of TP53 phosphorylation; the level of total TP53 remained unchanged. ONG41008 also induced expression of p21 and p16 in a concentration-dependent manner; this finding corroborated the results of ICC experiments. Next, we treated A549 cells with TGFβ to increase proliferation in the presence or absence of ONG41008. A significant proportion of A549 cells died after 6 days of treatment with ONG41008 and all cells died after 15 days of treatment, whereas control A549 cells remained alive (Figure 7B).

**Figure 7.**
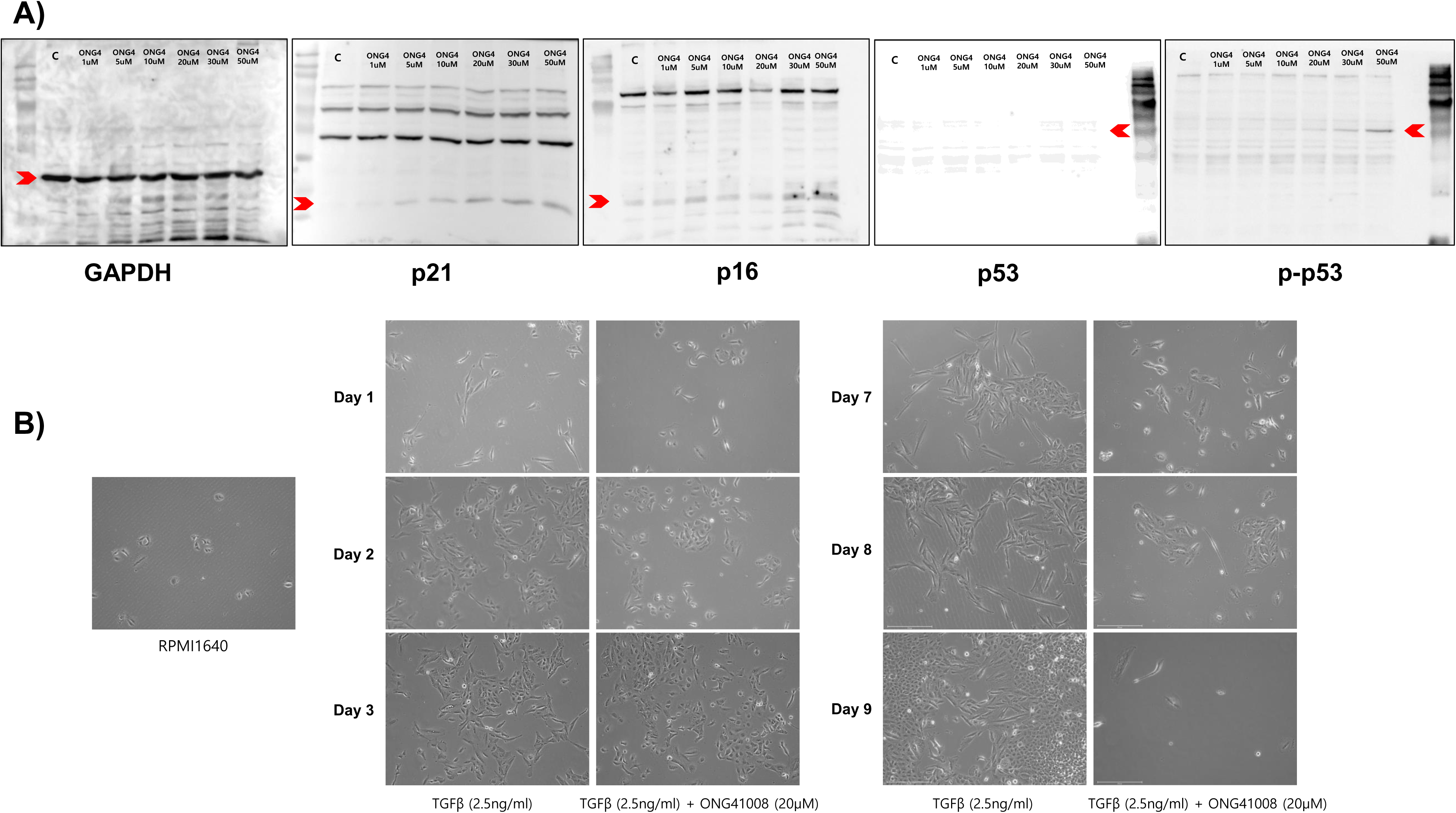
Western blot analysis of TP53, p21, and p16 expression and microscopic observation of A549 cell death. A) Cell lysates were prepared from A549 cells treated with 1 μM to 50 μM ONG41008 for 24 hr and subjected to western blot using the indicated antibodies. Red arrows represent the molecular mass of the respective proteins. B) A549 cells were continually treated with 20 μM ONG41008 for 15 days in the presence of TGFβ (2.5 ng/mL) to maintain cell viability. RPMI media was used as a control. Cell death was monitored using phase-contrast microscopy.

To further explore the senolysis induced by ONG41008, PANC1 cells were employed and treated with ONG41008 and cell viability was measured using a CCK-8 assay. The IC_50_ value was 11.3 μM (Figure 8A). To verify if cellular senescence or senolysis occurred, western blot analyses were conducted, and the results are shown in Figure 8B. Induction of p-TP53, p-Rb, and caspase-3 expression and cleavage of PARP were evident, suggesting that ONG41008 induced senolysis in PANC1 cells. Furthermore, ICC experiments showed that induction of p-TP53 expression and translocation to nucleus occurred (Figure 8C). We were able to show by real-time live imaging that ONG41008 efficiently induced a potent cytostatic effect on PANC1, human aggressive ductal adenocarcinoma within 24hr followed by rapidly killing PANC1(Supplementary Video II).

**Figure 8.**
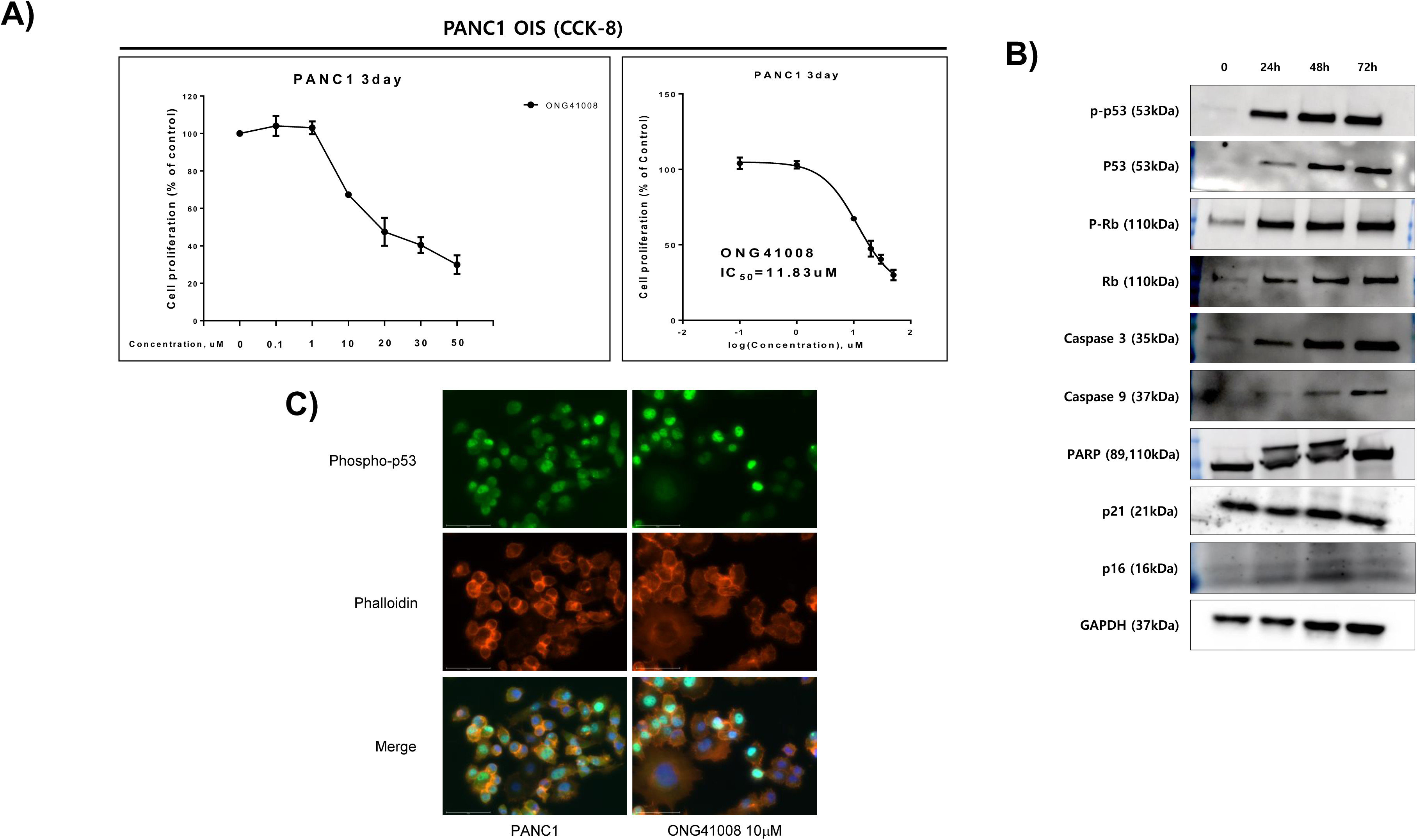
Oncogene-induced senescence by ONG41008 in human pancreatic cancer cells (PANC1). A) PANC1 cells were stimulated with 0.1 μM to 50 μM ONG41008 for 3 days. The cell survival rate was measured using a CCK-8 assay, and the IC_50_ value of ONG41008 was calculated on day 3. B) PANC1 cells were treated with 20μM ONG41008 for 24hr, 48hr and 72hr. Western blot analysis was conducted to detect p53, phospho-p53, Rb, phospho-Rb, caspase3, caspase9, PARP, p21, p16, and GAPDH. C) PANC1 cells were stimulated with 10 μM ONG41008 for 6 hr. ICC was conducted with phospho-p53 (green), phalloidin (red), and DAPI (blue).

Taken together, these results show that ONG41008 induced cellular senescence in cancer cells followed by senolysis.

### Recognition of the intracellular microenvironment by ONG41008

As described above, ONG41008 induced cellular senescence in DHLF but not in NHLF. Especially, ONG41008 did not induce apoptosis in DHLF. However, nintedanib induced robust apoptosis in DHLF, prompting us to hypothesize that ONG41008 might affect pathogenic or aged-cell microenvironments but not affect homeostatic intracellular microenvironments. This hypothesis is very challenging to test. First, we attempted to confirm if NHLF are unresponsive to ONG41008 but undergo rapid cellular toxicity in response to cisplatin or nintedanib. As shown in Figures 9A and 9B, ONG41008 exerted no apoptotic effects on NHLF and it was not possible to obtain an IC_50_ value, whereas cisplatin and nintedanib killed NHLF with IC_50_ values of 18.14 μM and 4.59 μM, respectively. We tested this hypothesis in human primary prostate epithelial cells (HPrECs). |In these cells, the IC_50_ value of ONG41008 was 46.77 μM and the IC_50_ value of cisplatin was IC_50_=0.14 μM, indicating that ONG41008 did not much impact HPrEC in terms of apoptosis (Figure 9C and 9D).

**Figure 9.**
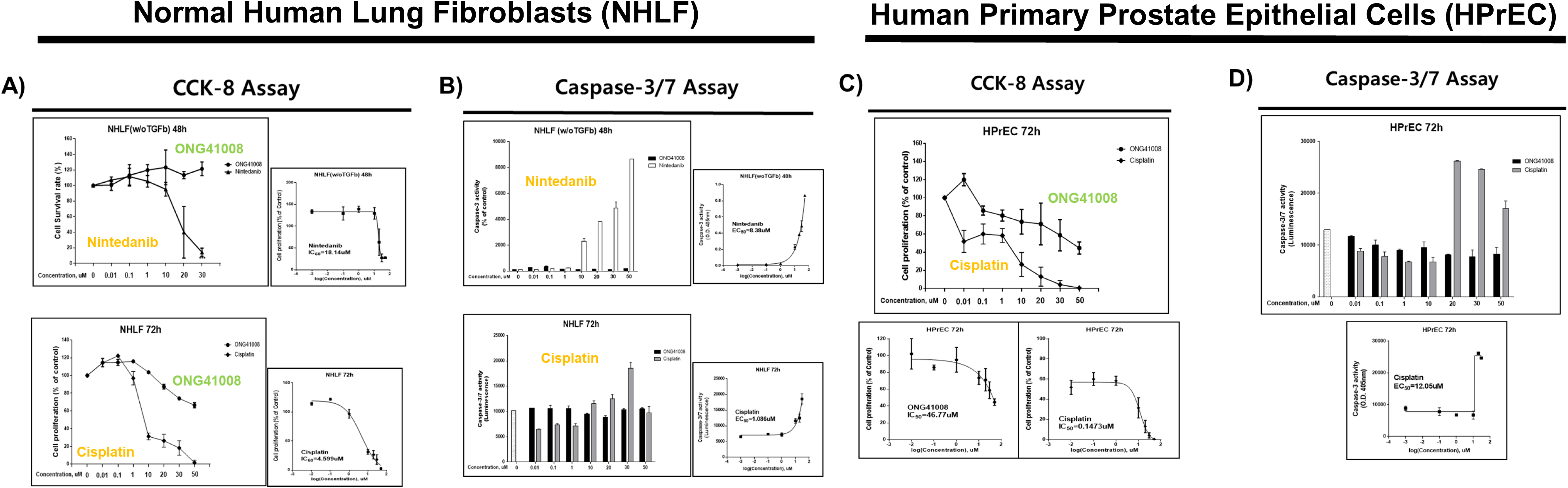
A comparison of the apoptotic effects of ONG41008, nintedanib, and cisplatin and the survival rate of NHLF and HPrEC. A) NHLF were stimulated with ONG41008 or nintedanib for 48 hr, and stimulated with ONG41008 or cisplatin for 72 hr. The cell survival rate was measured using a CCK-8 assay. The IC_50_ values of nintedanib and cisplatin were calculated. B) NHLF were stimulated with ONG41008 or nintedanib for 48hr and stimulated with ONG41008 or cisplatin for 72 hr. An activated caspase-3 assay was performed, and the EC_50_ values of nintedanib and cisplatin were calculated. C) Human prostate epithelial cells (HPrECs) were stimulated with ONG41008 or cisplatin for 72 hr. The cell survival rate was measured using a CCK-8 assay. The IC_50_ values of ONG41008 and cisplatin were calculated. D) HPrECs were stimulated with ONG41008 or cisplatin for 72 hr. An activated caspase-3 assay was performed, and the EC_50_ value of cisplatin was calculated with a sigmoidal, four-parameter logistic in GraphPad Prism version 7.00.

Taken together, these results indicate that unlike cisplatin, ONG41008 has apoptotic effects on both A549 and PANC1 cells, and induces moderate senolysis, which eventually results in cell death. However, normal cells did not appear to be affected by ONG41008.

### ONG41008 induces cell-cycle arrest at G2/M and increases the NAD/NADH ratio

The involvement of TP53, p16, and p21 in the action of ONG41008 strongly indicates that ONG41008 plays an important role in controlling the cell cycle. We conducted cell-cycle analysis using PI staining 24 hr and 48 hr after treatment of A549 cells with ONG41998 or fisetin as a control. ONG41008 as well as fisetin induced cell-cycle arrest at the G2/M stage (Figure 10A). Abrogation of CDK2 and CDK6 activity by ONG41008 could be further evidence for G2/M arrest. Metabolic regulation is a key driver of cell fate, determining if the cell cycle is controlled or whether cells undergo apoptosis (32). It is well-established that the NAD/NADH ratio plays a central role in regulating energy metabolism, including the control of glycolysis and the TCA cycle, and substantially affects mitochondrial functions such as those involved in disease pathogenesis or aging (33). We hypothesized that ONG41008 could influence the NAD/NADH ratio, based on the hitherto observation that ONG41008 could ‘sense’ the intracellular microenvironment and so distinguish normal cells from those undergoing uncontrolled proliferation, such as tumor cells or pathogenic myofibroblasts. When A549 cells were stimulated with ONG41008, the NAD/NADH ratio increased to between 60–80. (Figure 10B). However, SAHA did not affect the NAD/NADH ratio. In the absence of ONG41008, the NAD/NADH ratio in A549 cells had a negative value, suggesting that A549 cells may exclusively utilize anaerobic glycolysis when they are exposed to ONG41008.

**Figure 10.**
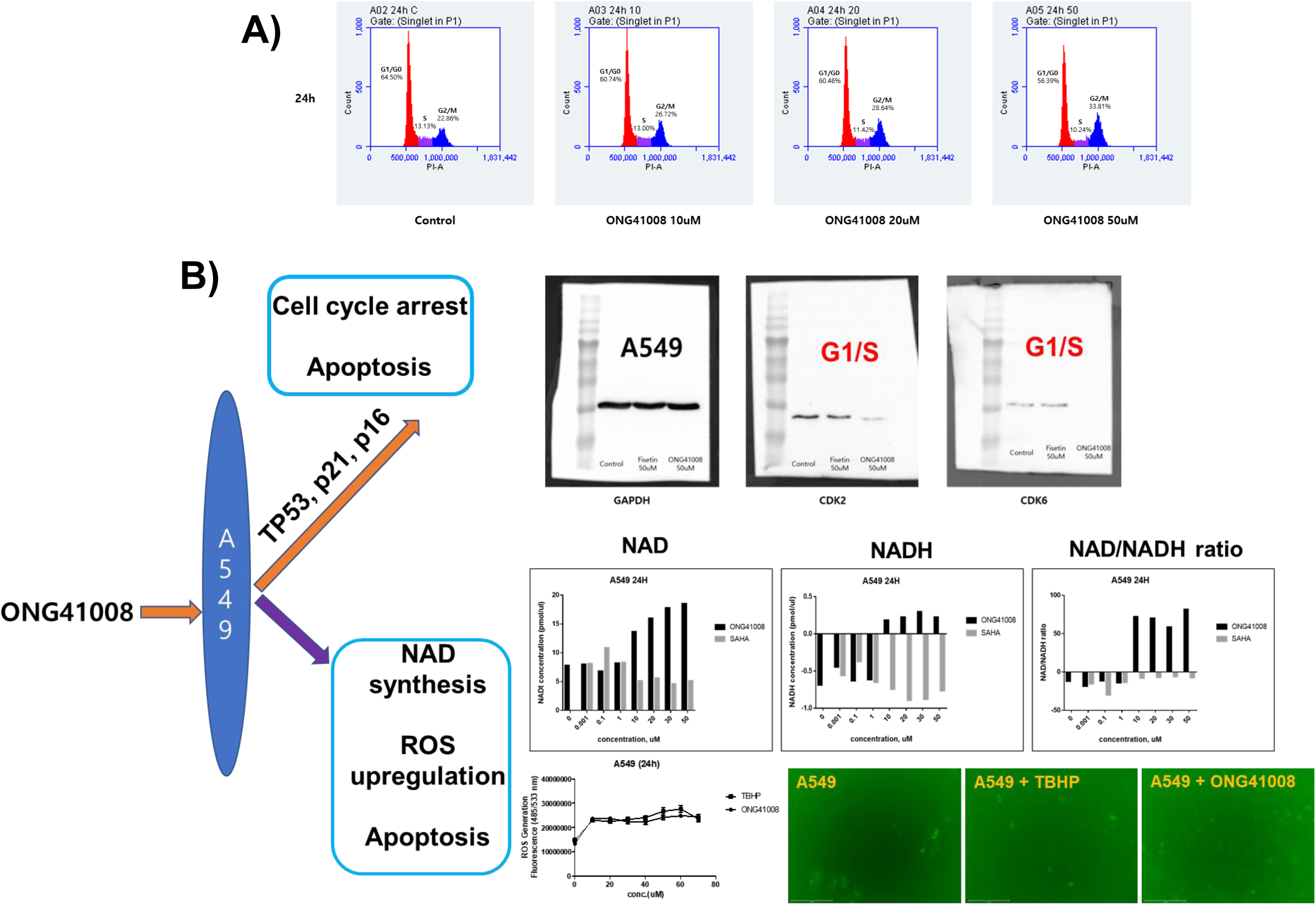
ONG41008-mediated cell-cycle arrest and induction of NAD/NADH and ROS expression. A549 cells were stimulated with 50 μM fisetin or ONG41008 for 24 hr, then cell lysates were collected. CDK2 and CDK6 expression was detected by western blot analysis. A549 cells were stimulated with medium alone, varying concentrations of SAHA or ONG41008 for 24 hr. Cell lysates were prepared and the NAD/NADH ratio was measured. The intracellular ROS levels in A549 cells was examined using H2DCFDA staining and fluorescence spectroscopy and fluorescence microscopy. TBHP was used as a positive control

## Discussion

Fibrosis and cancer are often intractable and fatal diseases with a pathogenesis that is directly related to cell-cycle regulation. Under normal conditions, the cell cycle is meticulously controlled such that proliferation, differentiation, and apoptosis are coordinated. Dysfunction of this tight regulation, which leads to uncontrolled proliferation, may increase the likelihood of serious diseases. Currently used anti-fibrotic and anti-cancer drugs largely target uncontrolled proliferation (34). Yet such drugs are associated with several side effects. Immune checkpoint inhibitors, which enable the host immune system to destroy cancer cells, are becoming a very successful anti-cancer strategy and are associated with fewer side effects than drugs that target proliferation (35). Here, we show that RS (senescence) and OIS (senolysis) induced by ONG41008 caused the proliferative arrest of DHLF and A549 cells, thereby preventing uncontrolled proliferation. Although we have not tested the effects of ONG41008 in many normal (that is non-cancer and non-fibrotic) cell lines, the compound was clearly able to distinguish pathogenic myofibroblasts and several cancer cells from normal lung fibroblasts and primary prostate epithelial cells.

Pathogenic myofibroblasts and cancer cells both gain ATP from aerobic glycolysis, resulting in a dysregulated redox potential that manifests as alterations in the NAD/NADH ratio, ATP levels, or intracellular pH. The CS is found in flavones or isoflavones (36), and captures UV to protect plants from UV damage. Although over 10,000 CS derivatives have been identified, only a limited number have anti-fibrotic effects. This finding suggests that in addition to the CS, a variety of side chains linked to the CS are needed for the compound to have anti-fibrotic effects. Another study reported that the CS contained with the structure of SB203580 is important in mediating the anti-cancer activity of this compound (37). Since ONG41008 can distinguish pathogenic myofibroblasts (DHLF) and cancer cells (A549, PANC1, MCF7, and PC3 cells) from normal cells, we propose that the CS might recognize features of the intracellular microenvironment such as NAD, NADH, pH, or ATP that act as energy sensors. As ONG41008 normalized the NAD/NADH ratio to a value of 60–100, we speculate that the normalized NAD/NADH ratio may influence ROS production in ONG41008-treated A549 cells, thereby partly contributing to senolysis. Because ONG41008 increases intracellular NAD concentrations, it is also possible that the NAD rate-limiting enzyme, NAMPT (nicotinamide phosphoribosyltransferase), is the molecular target of ONG41008.

Overall, our findings indicate that ONG41008 is a potent inducer of cellular senescence (RS and OIS) that arrests the pathogenic proliferation of myofibroblasts or cancer cells, thus preventing the uncontrolled proliferation that occurs in diseased states.

## Acknowledgments

We are thankful to all the Osteoneurogen researchers and administrative workers who have helped us and made the current manuscript possible. This study was supported by an intramural fund from Osteoneurogen, Inc. (ONG-400021).

## Competing financial interests

B-S Youn, I-H Kim, and H-S Kim retain shares in Osteoneurogen, M-K Meang retains a stock option, and SB Kim is employed by Osteoneurogen. The current study presented in this manuscript is the subject of a Korean provisionary patent.

## Author contributions

B-S Youn conceived the idea and wrote the manuscript. MK Maeng conducted the majority of the experimental procedures. SB Kim conducted experiments that determined the effect of the compound on fibroblasts and cancer cells, leading us to scrutinize cellular senescence. I-H Kim and H-S Kim continued to support for provision of various cancer cell lines and cell cycle analysis and to establish a theme termed Intracellular Molecular Recognition (IRM) together with us.

## *Footnotes

The following abbreviated terms were mainly used in this manuscript;

CS: Chromone Scaffold
CSD: Chromone Scaffold Derivatives
DHLF: Diseased Human Lung Fibroblasts from IPF patients
NHLF: Normal Human Lung Fibroblasts
RS: Replicative Senescence
OIS: Oncogene-Induced Senescence
MNC: Multinucleation
MTMP: Mitochondrial Membrane Potential
NAD: nicotinamide adenine dinucleotide
NAMPT: Nicotinamide Phosphoribosyltransferase

## Materials and Methods

### Cell culture and reagents

DHLF were purchased from Lonza (Basel, Switzerland) and cultured in fibroblast growth medium (FBM, Lonza, Walkersville, MD, USA). Recombinant human TGFβ and PDGF were obtained from Peprotech (Rocky Hill, CT, USA) and used at a final concentration of 5 ng/ml. Chemically synthesized ONG41008 was obtained from Syngene International Ltd. (Bangalore, India), dissolved at a stock concentration of 50 mM in DMSO, and stored in aliquots at -20°C. DMSO was used as a control. The RAW264.7 cell line was purchased from the Korean Cell Line Bank (Seoul, Korea) and cultured in RPMI supplemented with 10% FBS and 1% P/S (Welgene, Seoul, Korea). LPS was purchased from Sigma and used at a final concentration of 100 ng/ml.

### Immunocytochemistry

Cells were fixed using 4% paraformaldehyde, permeabilized with 0.4% TritonX100, blocked with 1% BSA and incubated with rhodamine phalloidin (Thermo Fisher, MA, USA), anti-GATA6 (Abcam, Cambridge, UK), anti-p53 (Cell Signaling Technology, Beverly, MA, USA), p21(Abcam, Cambridge, UK), p16-INK4A (Proteintech, IL, USA), ZEB1 (Cell Signaling Technology, Beverly, MA) for 4 hr at room temperature. After washing, cells were incubated with an Alexa Fluor 488 (Abcam, Cambridge, UK)-conjugated secondary antibody. Images were analyzed using EVOS M7000 (Invitrogen, CA, USA)

### Western blotting

A549 cells were seeded at 1 × 10^6^ cells/well in 100 mm cell culture dishes and incubated overnight, followed by treatment with various concentrations of ONG41008. After 24 hr, the cell lysates were clarified by centrifugation at 14,000 × g for 10 minutes and the supernatant was collected. The protein concentrations were quantified by the Bradford assay (Thermo Fisher, MA, USA). Thereafter, 25 µg of cellular protein was loaded on a 10% SDS-PAGE gel and transferred to nitrocellulose membranes. After blocking with 5% BSA, the membranes were incubated with anti-p53 (Cell Signaling Technology, Beverly, MA, USA), phosphor-p53 (Cell Signaling Technology, Beverly, MA, USA), p21 (Abcam, Cambridge, UK), p16-INK4A (Proteintech, IL, USA), and GAPDH (Abcam, Cambridge, UK) overnight at 4°C. After washing thoroughly, membranes were incubated with an HRP-conjugated secondary antibody. Protein bands were visualized using ECL reagent (Abfrontier, Korea) and Uvitec HD9 (Uvitec, Cambridge, UK).

### Live imaging

DHLF were seeded in 12-well cell culture plates and, after 24 hr, were treated with TGF beta (5 ng/ml), nintedanib (10 µM), pirfenidone (10 µM), and ONG41008 (10 µM). Cells were incubated in an EVOS M7000 CO_2_ incubation chamber (Invitrogen, CA, USA); cell morphology images were captured every 30 minutes. The collected images were assembled and are available as video (Supplementary Information).

### CCK-8 assay

Cells were seeded onto 96well plates at a density of 3×10^4^cells/well for 24hr. Then cells were treated with 0.01, 0.1, 1, 10, 20, 30 µM ONG41008, pirfenidone, or nintedanib. After treatment, the cell survival rate was measured using a CCK-8 assay (Dojindo, CK04) according to the manufacturer’s protocol.

### Caspase-3 assay

The caspase-3 activity was measured using a caspase-3 assay kit (Abcam, ab37401) following the manufacturer’s protocol. Cells treated with various concentrations of ONG41008, pirfenidone, or nintedanib were harvested and lysed on ice. The protein concentration was then measured by a BCA assay (ThermoFisher, 23227) and adjusted to 50 µg protein per 50 µl cell lysis buffer.

### Mitochondrial membrane potential assay

Cells were seeded in a 96-well plate for 24 hr, then exposed to different concentrations of ONG41008, pirfenidone, or nintedanib. The cells were then co-incubated with TMRE (Abcam, ab113852), for 30 min 37□ in the dark. Cells treated with 20 µM FCCP were used as a positive control. The mitochondrial membrane potential was measured following the manufacturer’s protocol.

### LDH assay

LDH release was detected by using an LDH assay kit (Abcam, ab56393). The cell culture plates were centrifuged at 480 g for 10 minutes, and supernatants (10 µl/well) were extracted into a new 96-well plate. Then, 100 µl of LDG reaction mix was added to each well and incubated for 30 minutes at room temperature. The absorbance values were measured at 450 nm on a microplate reader.

### NAD/NADH assay

Total NAD was extracted and quantified from A549 cell lysates using an NAD+/NADH colorimetric assay kit (Abcam, ab65348) following the manufacturer’s instructions. Cells (1×10^6^) were lysed in 400 µl NAD/NADH extraction buffer, filtered through a 10 kD spin column (ab93349) and measured neat or at a 1/5 dilution. Briefly, the amount of total NAD was calculated from a standard curve (pmol) divided by the sample volume added to the reaction well (μl) and multiplied by the dilution factor.

### RNA-seq, differential gene expression, and interactome analyses

Processed reads were mapped to the *Mus musculus* reference genome (Ensembl 77) using Tophat and Cufflink with default parameters. Differential analysis was performed using Cuffdiff using default parameters. Then, FPKM values from Cuffdiff were normalized and quantitated using the R Package tag count comparison (TCC) to determine statistical significance (e.g., p-values) and differential expression (e.g., fold-changes). Gene expression values were plotted in various ways (i.e., Scatter, MA, and Volcano plots) using fold-change values and an R-script developed in-house. The protein interaction transfer procedure was performed using the STRING database with the differentially expressed genes. A 60 Gb sequence was generated, and 10,020 transcripts were read and compared. The highest confidence interaction score (0.9) was applied from the *Mus musculus* reference genome, and information about interactions was obtained based on text mining, experiments, and databases (http://www.string-db.org/). Due to proprietary company information, we have not provided a detailed interpretation of the RNA-Seq or interactome data, but we have provided sufficient analysis to support our assertions in Results and Discussion.

### Reverse transcriptase PCR and real-time PCR

Cells cultured in either 12- or 24-well plates were washed twice with cold PBS and harvested using a TaKaRa MiniBEST universal RNA extraction kit (Takara, Japan). RNA was purified using the same kit according to the manufacturer’s protocol. RNA was reverse-transcribed using a cDNA synthesis kit (PCRBio Systems, London, UK). Synthesized cDNA was amplified with StepOne Plus (Applied Biosystems, Life Technologies) and 2× qPCRBio probe mix Hi-ROX (PCRBio). Comparisons between mRNA levels were performed using the ΔΔCt method, with GAPDH as the internal control.

## Supplemental Information

**Suppl. Video I:** A video of high-density DHLF treated with various drugs and compounds for 72 hr; one image was taken every 30 minutes.

**Suppl. Video II:** PANC1 cells were cultured in CO2 incubation chamber connected with microscope for 6 days. The cells were stimulated with 20μM ONG41008 for 24hr, 72hr, or 96hr and wash with DPBS and continued to be cultured in plain medium till 6 days. Live imaging was acquired for comparison.

**Suppl. Table I**: Top-ranked replicative senescence-initiating genes: soluble factors / receptors. RNA-seq analysis was conducted using total RNAs prepared from DHLF or DHLF treated with ONG41008 for 24 hr. The 20 top-ranked genes showing p>0.005 were selected.

**Suppl. Figure 1: ONG41008 and ONG21001 induce cellular senescence leading to multinucleation** A549 cells were stimulated with 20 μM ONG41008, ONG21001, hispidulin, jaceosidin, or apigenin for 72 hr and subjected to ICC using DAPI, and phalloidin. MNC are denoted with white circles.

**Suppl. Figure 2 and Figure 3:** An RNA-Seq analysis was performed as described in the Materials and Methods. The genes were sorted so that three sets of genes with p-values of between 0.005 and 0.05, or between 0.05 and 0.5 were selected; the selected genes were further analyzed using the STRING program, which resulted in three interactomes.

**Suppl. Figure 4: Comparison of senogenicity** A549 were stimulated with ONG41008, Quercetin, Fisetin, or Dasatinib for 48hr and stained with phallodine and DAPI. Cell flatness was observed through phase-contrast microscopy. Multinucleation was highlighted with white circles.

**Suppl. Figure 5: ONG41008 induces cellular senescence in human aggressive cancer cell lines** A549, MCF7, PC3, or PANC1 cells was stimulated with 20 μM ONG41008 for 24 hr. ICC was conducted using phalloidin imaging. MNC are denoted with white circles.

**Suppl. Figure 6: Translocation of TP53 to the nucleus, and induction and nuclear translocation of p21 and p16 in ONG41008-treated A549 cells** A549 cells were treated with 1 μM to 10 μM of ONG41008 or control. ICC was conducted with anti-human TP53, anti-p21, anti-p16, or phalloidin in conjunction with DAPI. A) Translocation of p53 to the nucleus. B) Induction and translocation of p21 to the nucleus. C) Induction and translocation of p16 to the nucleus.

**Suppl. Figure 7: Translocation and accumulation of p16 to nucleus ONG41008-treated A549 cells** A549 cells were treated with 20 μM of ONG41008 or control (RPMI medium). ICC was conducted with anti-p16 or phalloidin in conjunction with DAPI.

## Notes

**Funding:** funded by an Osteoneurogen intramural fund

### Summary of Updates

bioRxiv Dear Editor, We would like to revise the existing preprint entitled "A small molecule that promotes cellular senescence prevents fibrogenesis and tumorigenesis in vitro"(bioRxiv 2021.06.01.446522; doi: https://doi.org/10.1101/2021.06.01.446522). The major reason is we have enormously fortified the preprint with cellular senescence and oncogene-induced senescence (OIS) data associated with our drug candidate called ONG41008 with depth and width. Several senolytic drugs were included and compared with ONG41008 (Figures 3, 4, and 5 are new data. Running title has been changed: Discovery of a senotherapeutic molecule. We believe that you shall find much-evolving data by reading the revised manuscript. We look forward to hearing from your reply. Sincerely, Byung-Soo Youn, Ph.D. CEO (Founder) #411, 128, Gasan digital 1-ro, Geumcheon-gu, Seoul, 08507 Republic of Korea Tel) 822- 6267-2739 Fax) 822-6267-2740 Cell)82-2- 10-2793-2737 Web) www.osteoneurogen.com

