## supplementary figures for "A small molecule that promotes cellular senescence prevents fibrogenesis and tumorigenesis *in vitro*"

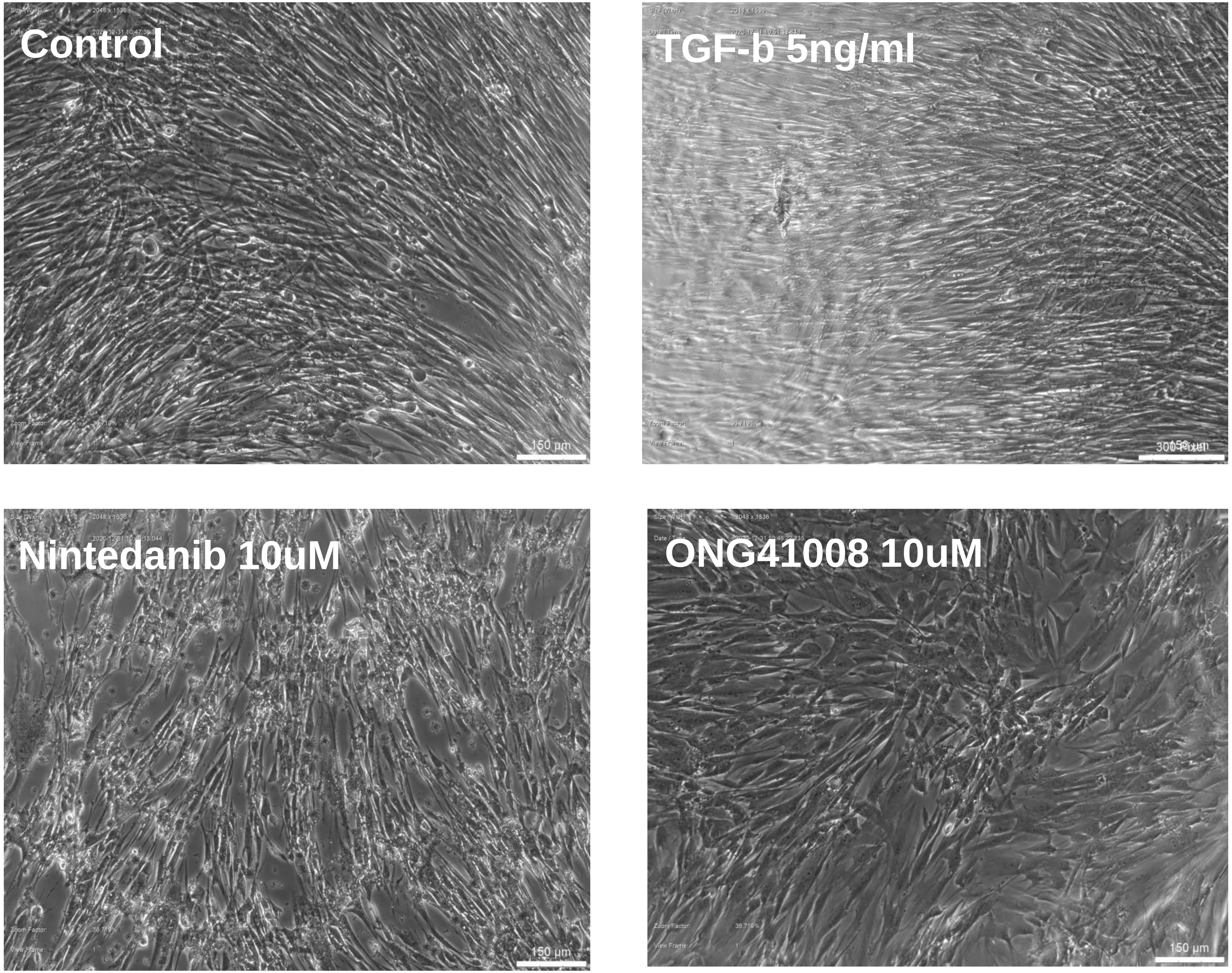


**Supplementary Video I**

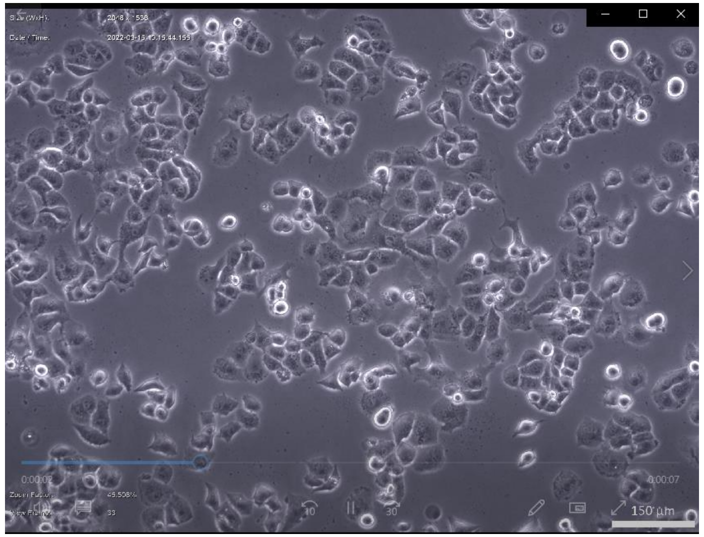


PANC1

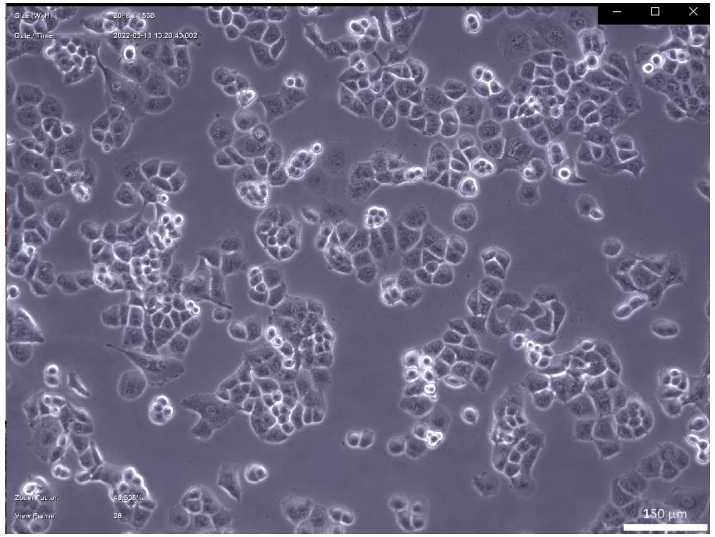


ONG41008 20μM (24hr)

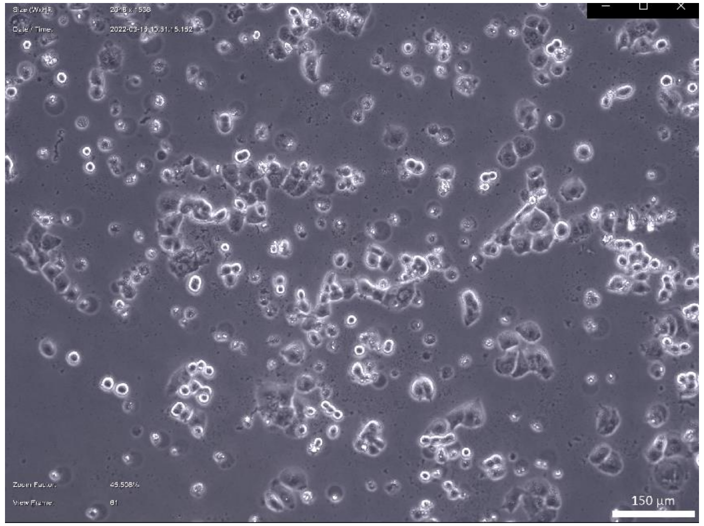


ONG41008 20μM (72hr)

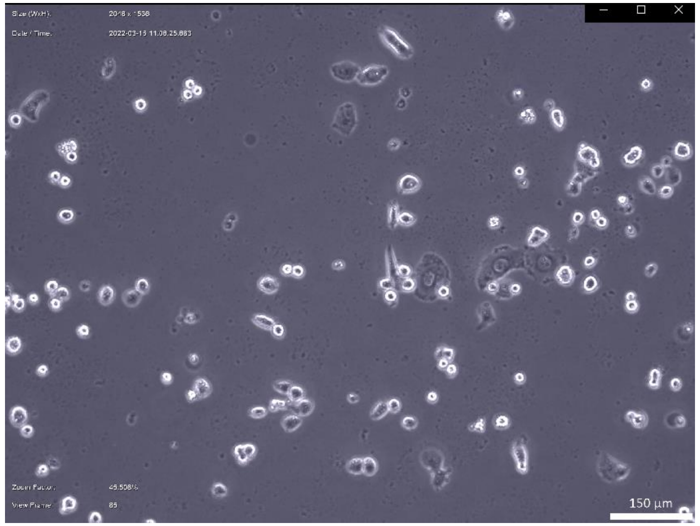


ONG41008 20μM (96hr)

**Supplementary Video II**

| GeneName | Desc | log2fc | p-value | q-value |
|---|---|---|---|---|
| **NRP1** | neuropilin 1 | -2.78 | 5.00E-05 | 0.0159 |
| **MMP1** | matrix metallopeptidase 1 (interstitial collagenase) | -3.12 | 5.00E-05 | 0.0159 |
| **FARP1** | FERM, RhoGEF (ARHGEF) and pleckstrin domain protein 1 (chondrocyte-derived) | -2.53 | 5.00E-05 | 0.0159 |
| **THSD4** | thrombospondin, type I, domain containing 4 | -4.56 | 5.00E-05 | 0.0159 |
| **NPTX1** | neuronal pentraxin I | -3.84 | 5.00E-05 | 0.0159 |
| **SLC14A1** | solute carrier family 14 (urea transporter), member 1 (Kidd blood group) | -4.74 | 5.00E-05 | 0.0159 |
| **CCBE1** | collagen and calcium binding EGF domains 1 | -3.14 | 5.00E-05 | 0.0159 |
| **SPPL2B** | signal peptide peptidase like 2B | | 5.00E-05 | 0.0159 |
| **TBC1D5** | TBC1 domain family, member 5 | -3.66 | 5.00E-05 | 0.0159 |
| **ABI3BP** | ABI family, member 3 (NESH) binding protein | -2.92 | 5.00E-05 | 0.0159 |
| **PTX3** | pentraxin 3, long | -4.05 | 5.00E-05 | 0.0159 |
| **LPP** | LIM domain containing preferred translocation partner in lipoma | -3.72 | 5.00E-05 | 0.0159 |
| **SCFD2** | sec1 family domain containing 2 | -3.73 | 5.00E-05 | 0.0159 |
| **MAPK10** | mitogen-activated protein kinase 10 | -3.12 | 5.00E-05 | 0.0159 |
| **CAMK2D** | calcium/calmodulin-dependent protein kinase II delta | -3.02 | 5.00E-05 | 0.0159 |
| **RPS6KA2** | ribosomal protein S6 kinase, 90kDa, polypeptide 2 | -2.77 | 5.00E-05 | 0.0159 |
| **ANLN** | anillin, actin binding protein | -2.75 | 5.00E-05 | 0.0159 |
| **CACNA2D1** | calcium channel, voltage-dependent, alpha 2/delta subunit 1 | -3.13 | 5.00E-05 | 0.0159 |
| **EXOC4** | exocyst complex component 4 | -3.14 | 5.00E-05 | 0.0159 |
| **TNFRSF11B** | tumor necrosis factor receptor superfamily, member 11b | -2.72 | 5.00E-05 | 0.0159 |
| **PRUNE2** | prune homolog 2 (Drosophila) | -2.54 | 5.00E-05 | 0.0159 |

**Supple. Table I**

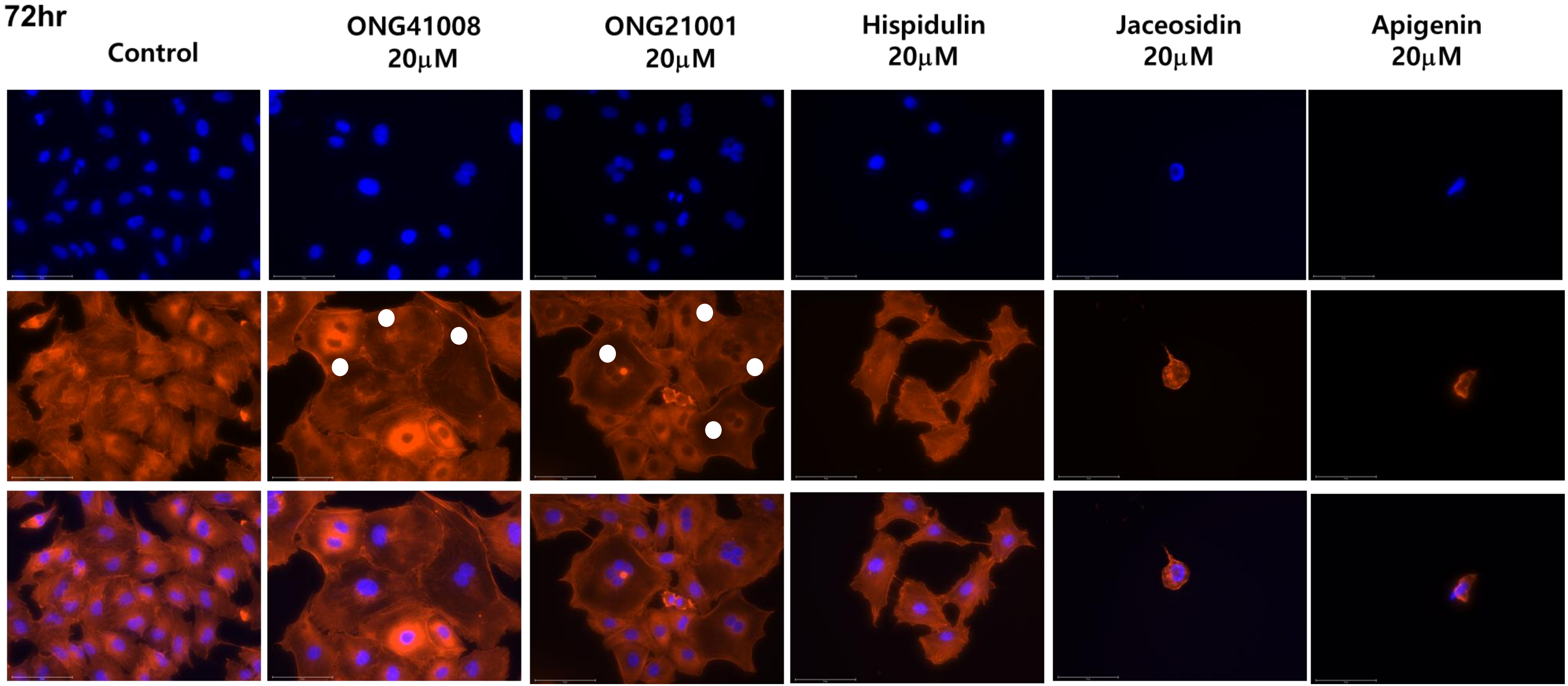


Supple. Figure 1

### Metabolic shift / Histone modification

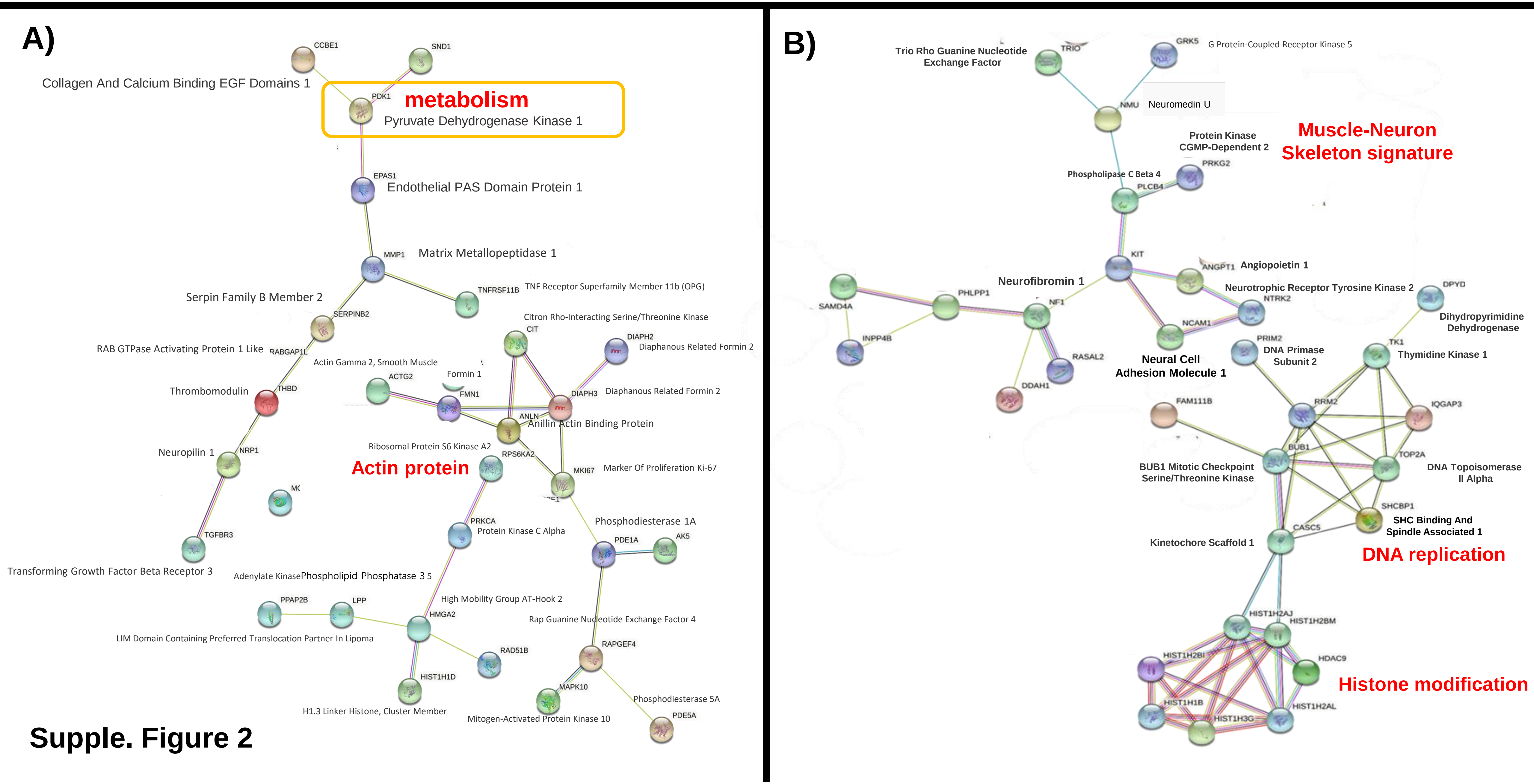


Supple. Figure 2

### DNA replication / Chromatin Remodeling / Cell Division Control

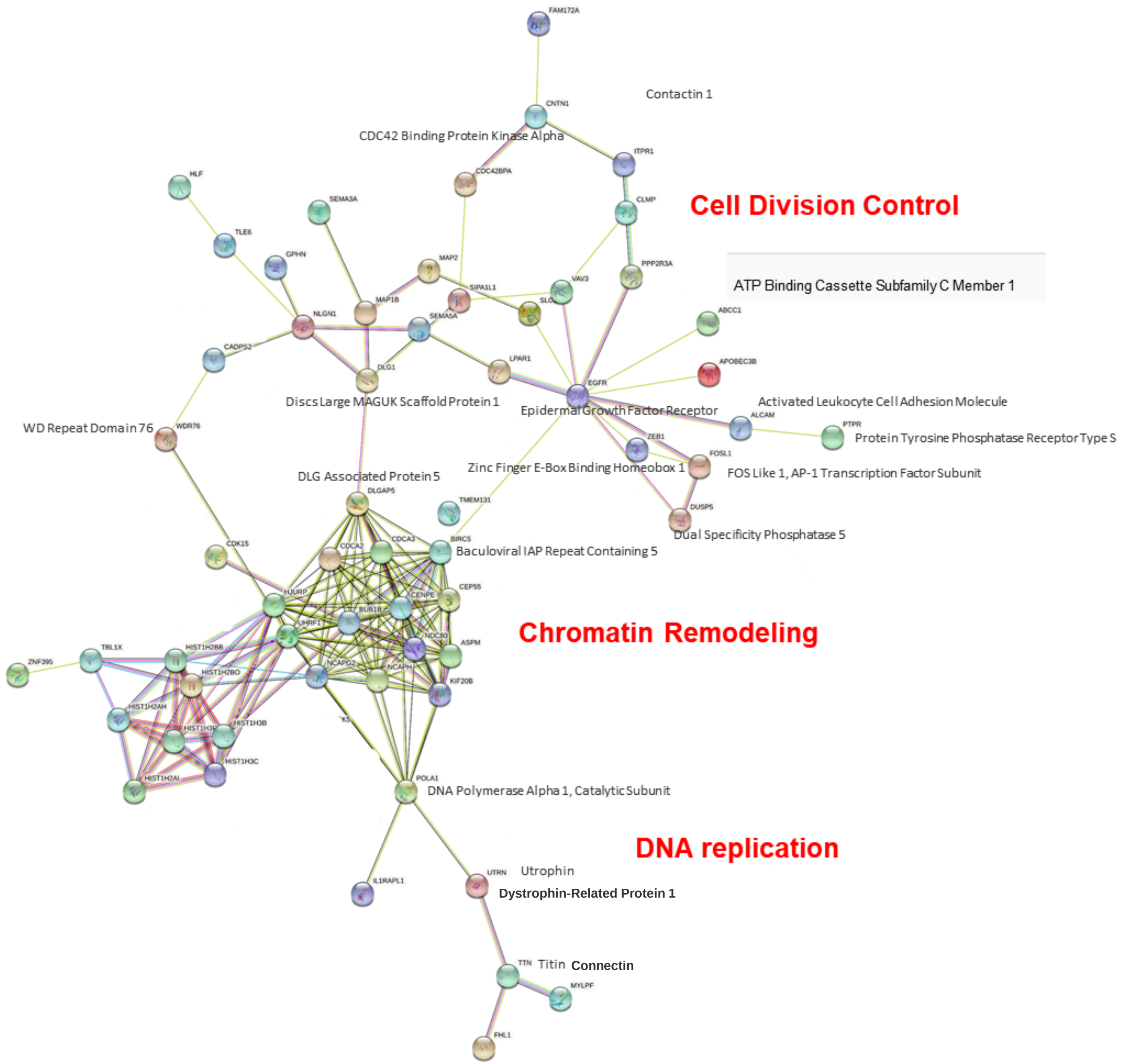


Supple. Figure 3

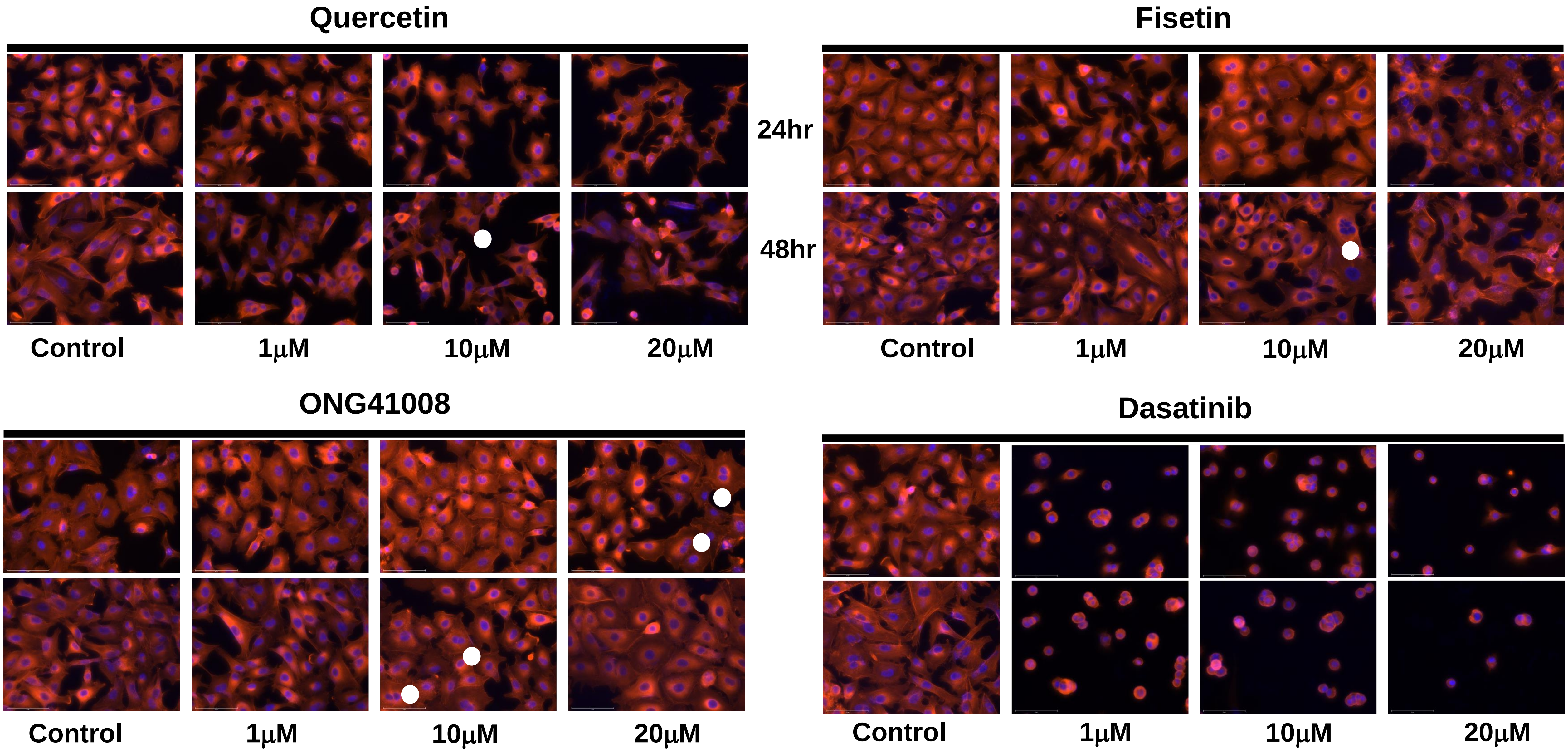


Supple. Figure 4

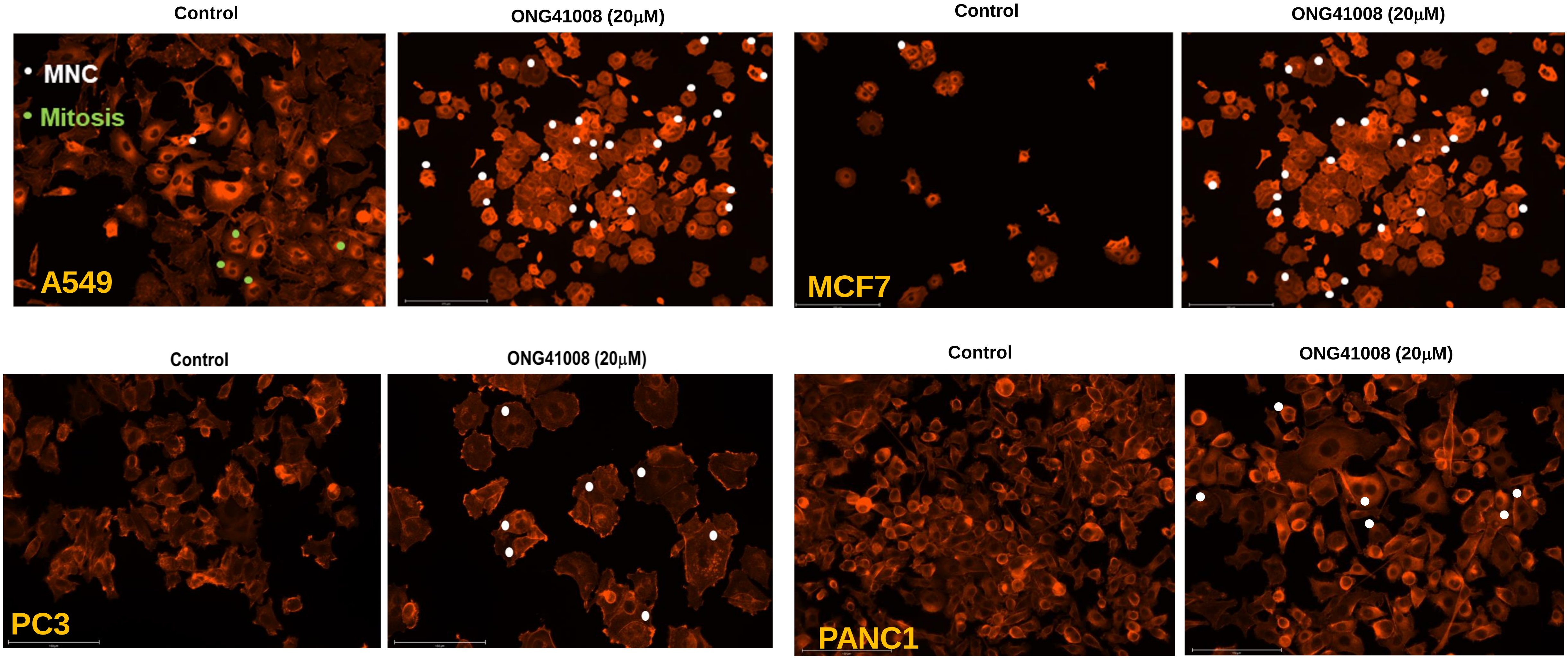


**Supple. Figure 5**

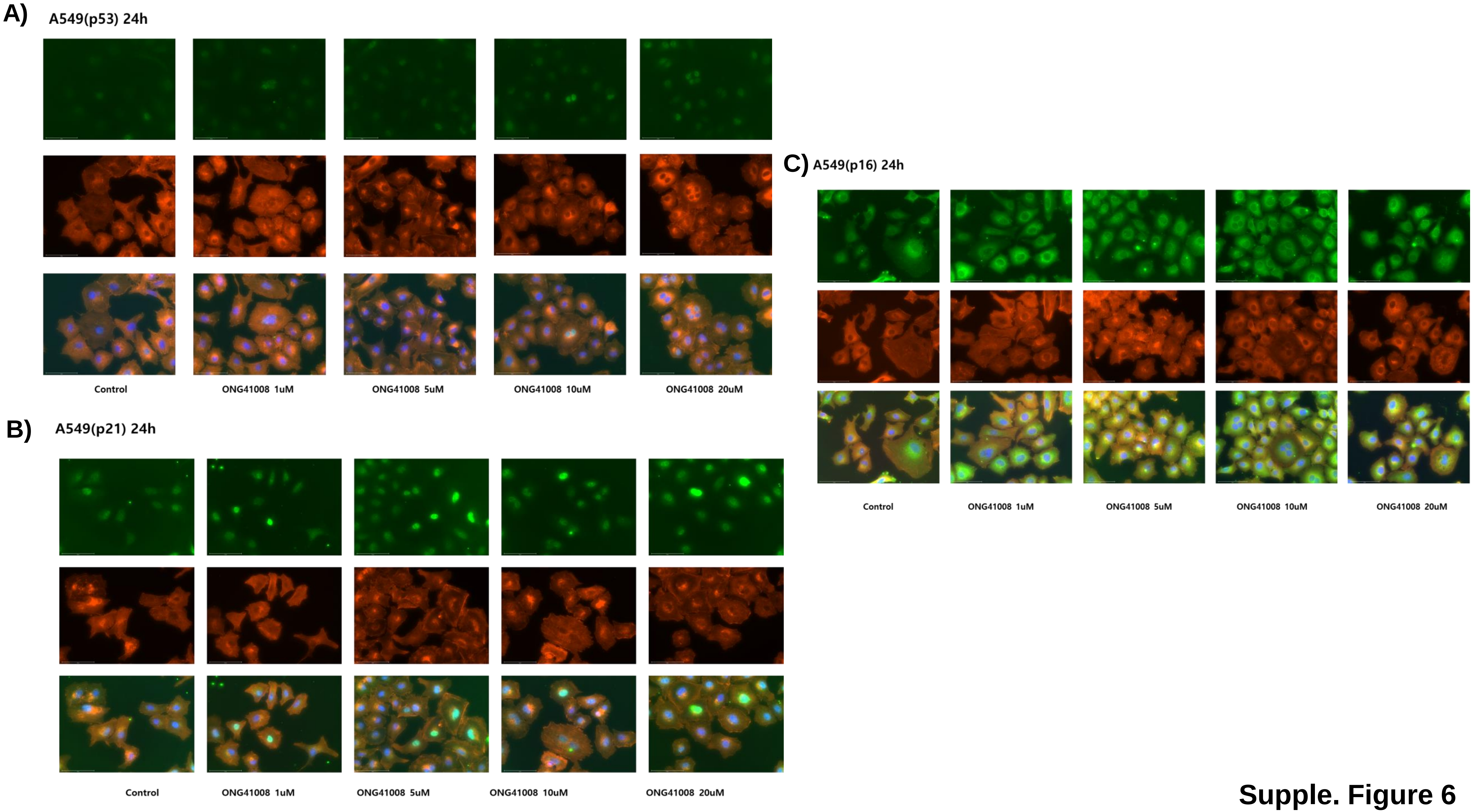


Supple. Figure 6

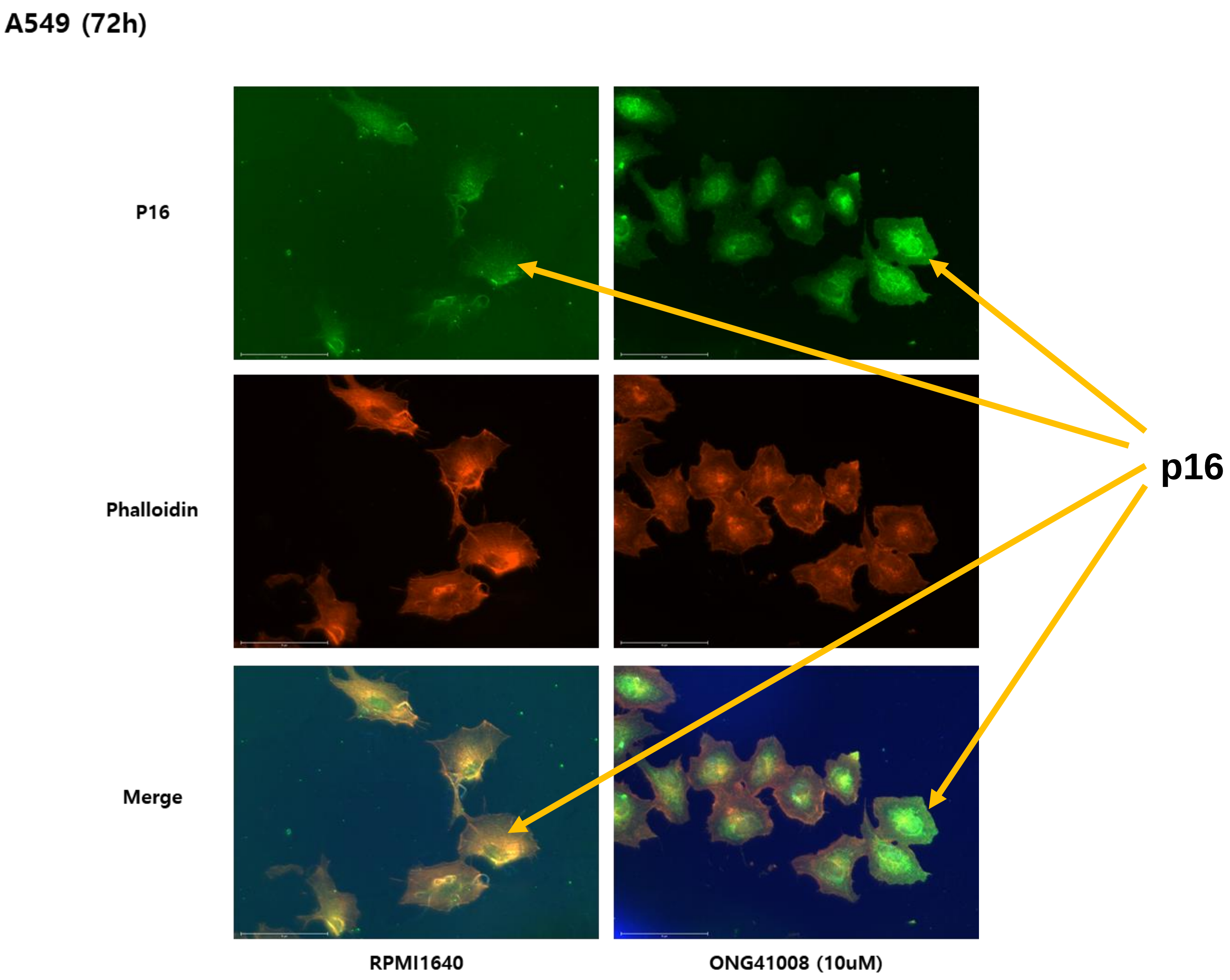


Supple. Figure 7
